# Probabilities of tree topologies with temporal constraints and diversification shifts

**DOI:** 10.1101/376756

**Authors:** Gilles Didier

## Abstract

Dating the tree of life is a task far more complicated than only determining the evolutionary relationships between species. It is therefore of interest to develop approaches apt to deal with undated phylogenetic trees.

The main result of this work is a method to compute probabilities of undated phylogenetic trees under Markovian diversification models by constraining some of the divergence times to belong to given time intervals and by allowing diversification shifts on certain clades. If the diversification models considered are lineage-homogeneous, the time complexity of this computation is quadratic with the number of species of the phylogenetic tree and linear with the number of temporal constraints.

The interest of this computation method is illustrated with three applications, namely,

- to compute the distribution of the divergence times of a tree topology with temporal constraints,
- to directly sample the divergence times of a tree topology, and
- to test for a diversification shift at a given clade.

## 1 Introduction

Estimating divergence times (i.e., the times of the speciation events corresponding to the internal nodes of a phylogenetic tree) is an essential and difficult stage of phylogenetic inference [24, 25, 18, 5, 22]. In order to perform this estimation, current approaches use stochastic models for combining different types of information: molecular and/or morphological data, fossil calibrations, evolutionary assumptions etc [35, 26, 9, 14]. An important point here is that dating speciation events is far more complicated and requires stronger assumptions on the evolutionary process than just determining the evolutionary relationships between species, not to mention the uncertainty with which divergence times can be estimated. It is therefore preferable to use, as much as possible, methods that do not require the exact knowledge of the divergence times. This is in particular true for studying questions related to the diversification process since the diversification process and divergence times are intricately linked. Diversification models are used in order to provide “prior” probability distributions of divergence times (i.e., which does not take into account information about genotype or phenotype of species [37, 16, 14]) [6, 15, 37]. Conversely, estimating parameters of diversification models requires temporal information about phylogenies.

We consider here Markovian diversification models (i.e., with independence and memoryless properties) among which the birth-death-sampling model is arguably the simplest realistic model since it includes three important features shaping phylogenetic trees [38, 39]. Namely, it models cladogenesis and extinction of species by a birth-death process and takes account of the incompleteness of data by assuming a uniform sampling of extant taxa. The birth-death-sampling model has been further studied and is currently used for phylogenetic inference [31, 33, 15, 6]. Since assuming constant diversification rates along time is sometimes unrealistic, the birth-death-sampling model has been extended in various ways. The generalized birth-death process proposed in [17] enables us to consider time-varying rates. In the model presented in [32], the diversification rates are piecewise-constant and the model supports the sampling of lineages not only at the present time but also at given past times in order to model mass extinction events. We combined the features of these two models to devise the *sampled-generalized-birth-death model* which allows both times-varying rates and past and extant samplings (Appendix B). The main goal of this work is to present methods to compute probabilities of undated phylogenies under certain assumptions about divergence times and about the diversification process under general models. Though this study focuses on methodological and computational aspects, three applications illustrating its practical interest are provided.

The first result is a method to compute the probability, under a Markovian diversification model, of a tree topology in which the divergence times are not exactly known but can be “constrained” to belong to given time intervals. This computation is performed by splitting the tree topology into small parts involving the times of the temporal constraints, referred to as *patterns*, and by combining their probabilities in order to get the probability of the whole tree topology. If the diversification model is lineage-homogeneous, the total time complexity of this computation is quadratic with the size of the phylogeny (i.e., its total number of nodes) and linear with the total number of constraints. Its memory space complexity is quadratic with the size of the phylogeny. In practice, it can deal with phylogenetic trees with hundreds of tips on standard desktop computers.

This computation can be used to obtain the divergence time distributions of a given undated phylogeny with temporal constraints, which can be applied to various questions. First, it can be used for dating phylogenetic trees from their topology only, like the method implemented in the function *compute.brlen* of the R-package *APE* [12, 23]. It also enables us to visualize the effects of the model parameters on the prior divergence times distributions, to investigate consequences of evolutionary assumptions etc. Last, it can provide prior distributions in phylogenetic inference frameworks. Note that the ability to take into account temporal constraints on the divergence times is particularly interesting in this context since in the calibration process, fossil ages are generally used for bracketing some of the divergence times [20]. The computation of the divergence time distribution is illustrated with a contrived example in order to show the influence of the temporal constraints and on a real phylogenetic tree in order to show the influence of the parameters of a simple birth-death-sampling model on the divergence time distributions. A previous method for computing divergence time distributions under the birth-death model [10] is briefly recalled in Section 7.1. By reviewing this work, Amaury Lambert proposed an alternative method to compute the divergence time distribution with temporal constraints, which has the same computational complexity as the approach presented here. This alternative method is presented in Section 7.2. The methods presented in Sections 7.1 and 7.2 both require that the divergence times are independent and identically distributed under the diversification model considered, an assumption which is not necessary with the approach presented here.

The computation of the probability of a tree topology under a given model allows us to sample all its divergence times under this model. In particular, this sampling procedure can easily be integrated into phylogenetic inference software [8, 27], e.g., for proposing accurate MCMC moves.

A second result shows how to calculate the probability of a tree topology in which a given clade is assumed to diversify following a diversification model different from that of the rest of the phylogeny. A natural application of this computation is to test diversification shift in undated phylogenies. It is used to define a likelihood ratio test for diversification shift which is compared with three previous approaches studied in [36].

Last, the approach presented here can be extended in order to take into account fossils. In [4], we started to work in this direction by determining divergence time distributions from tree topologies and fossil ages under the fossilized-birth-death model in order to obtain better node-calibrations for phylogenetic inference.

C-source code of the software performing the computation of divergence time distributions and their sampling under (piecewise-constant-)birth-death-sampling model and the shift detection test is available at https://github.com/gilles-didier/DateBDS.

The rest of the paper is organized as follows. Diversification models and birth-death-sampling models are formally introduced in Section 2.1. Section 3 presents definitions and some results about tree topologies. The standard and special patterns, i.e., the subparts of the diversification process from which are computed our probabilities, are introduced in Section 4. Sections 5 and 6 describe the computation of the probabilities of tree topologies with temporal constraints and diversification shifts, and show that this computation is quadratic with the size of the tree topology. Divergence time distributions obtained on two examples are displayed and discussed in Section 7. The method for directly sampling the divergence times is described in Section 8. Last, Section 9 presents a likelihood ratio test derived from the computation devised here, for determining if a diversification shift occurred in a tree topology. Its accuracy is assessed and compared with three previous tests of diversification shift. Appendices start with a table of notations, followed with the presentation of the sampled-generalized-birth-death model then with the proofs of theorems.

## 2 Diversification models

The methods presented below apply to general diversification models. Namely, a diversification model Θ provides the parameters of a stochastic process which starts with a single lineage at time *s* and ends at time *e*, where *e* is usually the present time (both *s* and *e* are parameters of Θ). At any time *t* between *s* and *e*, two types of event may occur on a lineage alive at *t*: a *speciation* event, which gives rise to a new lineage and an *extinction* event which basically kills the lineage. We also assume that the lineages alive at the end time *e* are sampled in a certain way (Fig. 1). A lineage alive at the end time which is not sampled has to be interpreted as a taxa which is not included in the study, for instance because it is unknown.

**Figure 1:**
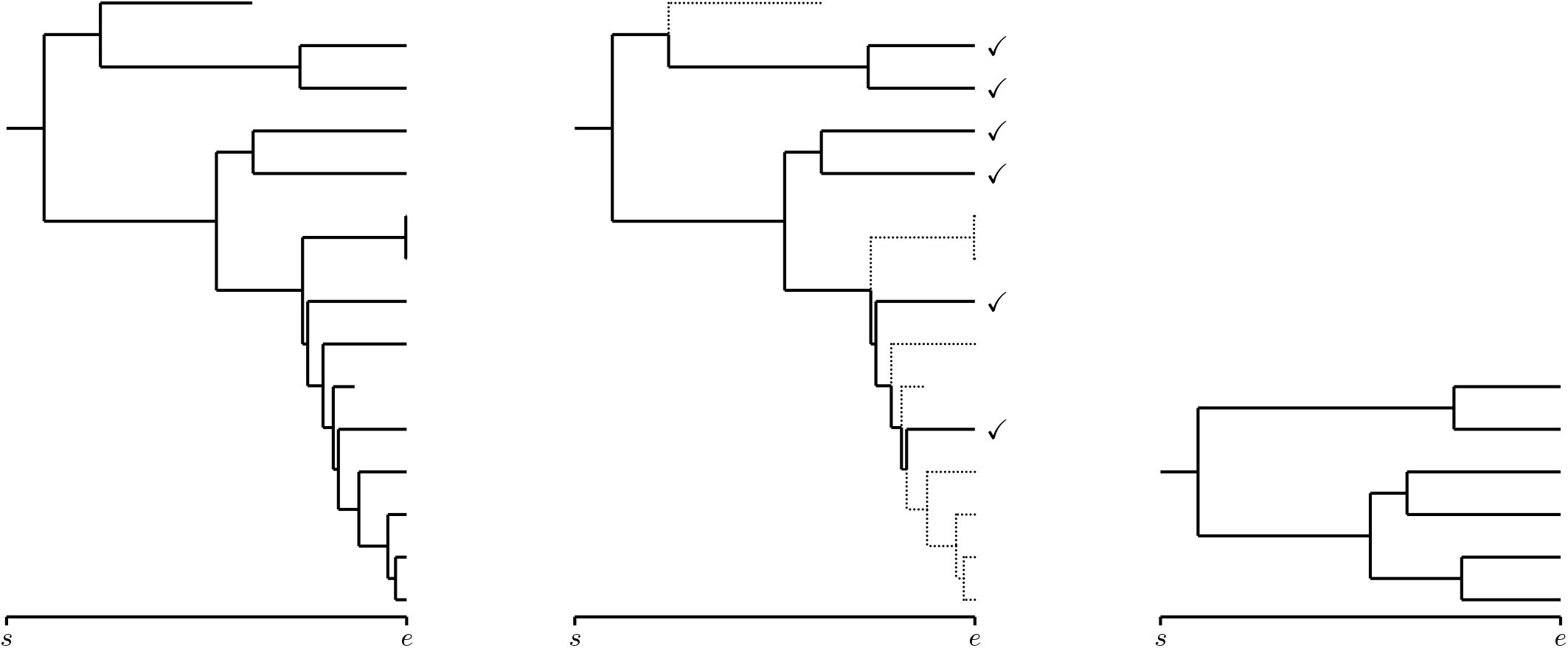
Left: the whole diversification process; Center: the part of the process that can be reconstructed is represented in plain – the dotted parts are lost (sampled extant species are those with ‘✓’); Right: the resulting phylogenetic tree.

An important point is to distinguish between the part of the process that actually happened, which will be referred to as the *complete process* (Fig. 1-Left) and the part that can be observed from the available information at the present time (i.e., from the sampled extant taxa), which will be referred to as the *reconstructed process* (Fig. 1-Right).

More formally, for all times *t* ∈ [*s, e*], a lineage alive at time *t* is *observable* if itself or at least one of its descendants are both alive and sampled at the end time *e*. We assume that the reconstructed process, which encompasses all the observable parts of the diversification, is the only information available.

Let Θ be a diversification model starting at *s* and ending at *e*, 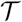 be a tree topology and *t* and *t*′ be two times such that *s* ≤ *t* ≤*t*′≤ *e*. We assume that we are able to compute under Θ:

- 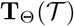, the probability that the reconstructed tree topology is 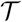 conditionally on the number of tips of 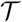,
- **Q**_Θ_(*t, t′, N*), the probability that a lineage alive at time *t* has *N* descendants at time *t*′ if *t′ < e* and *N* descendants *sampled* at time *e* otherwise,
- **O**_Θ_(*t*), the probability that a lineage alive at time *t* has at least a sampled descendant at the end time *e*.

For a diversification model starting at *s* and ending at *e* and a time *t* ∈ [*s, e*], we put Θ_[*t*]_ for the model Θ restricted to the time interval [*t, e*]. Namely, Θ_[*t*]_ models the evolution of a lineage alive at *t* until *e* under Θ.

A diversification model Θ is *Markovian* if conditionally of being alive at a time *t* between *s* and *e*, the evolution of a lineage from *t* is independent of its evolution before *t* and of that of the other lineages. A diversification model Θ is *lineage-homogeneous* if conditionally at occurring at a time *t*, any event of the process occurs on all the lineages alive at *t* with equal probabilities.

### 2.1 Birth-death-sampling models

Under a birth-death-sampling model, the dynamics of speciation and extinction of species follows a birth-death process with constant rates *λ* and *μ* both through time and lineage, starting at origin time *s* and ending at time *e* which is generally the present time [21]. Following [38], each extant species is assumed to be independently sampled at the end time *e* with probability *ρ*. The whole model will be referred to as the *birth-death-sampling model* and has thus five parameters:

- *s*: the origin time of the diversification process,
- *e*: the end/present time,
- *λ*: the speciation rate,
- *μ*: the extinction rate and
- *ρ*: the probability for an ending/extant taxa to be sampled.

Let us start by recalling some already derived probabilities of interest. By assuming that the diversification follows a simple birth-death process (i.e., with *ρ* = 1) with speciation rate *λ* and extinction rate *μ*, the probability *p_N_* (*t*) that a single lineage at time 0 has exactly *N* descendants at time *t* was given in [21]. We have that

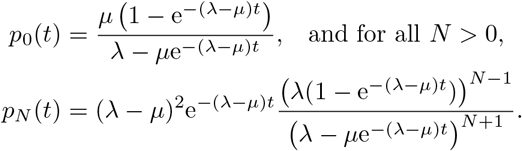

If one assumes that the diversification follows a birth-death-sampling process with speciation rate *λ* and extinction rate *μ*, the probability 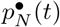 that a single lineage at time 0 has exactly *N* descendants sampled with probability *ρ* at time *t* was given in [38]. We have that

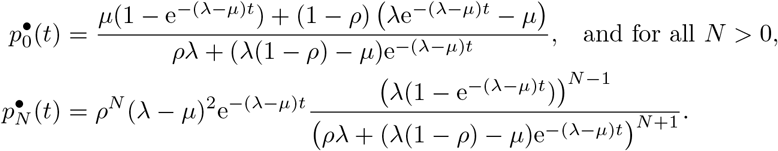

Let Θ = (*s, e, λ, μ, ρ*) be a birth-death-sampling model. For all pair of times *t* and *t*′ with *s ≤ t ≤ t′* and all number *N*, we define **Q**_Θ_(*t, t′, N*) as the probability under the model Θ that a lineage alive at time *t* has *N* descendants alive at time *t*′ if *t′ < e* and *N* descendants alive and sampled if *t*′ = *e*. We have that

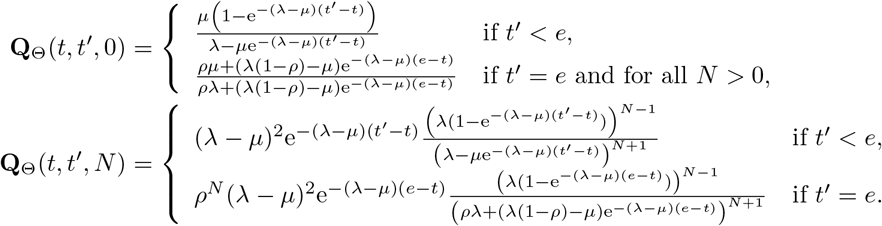

The probability **O**_Θ_(*t*) for a lineage living at time *t* in the complete diversification process (as in Figure 1-Left) to be observable (i.e., to be part of the reconstructed process) is the complementary probability of having no descendant sampled at time *e*. We have that

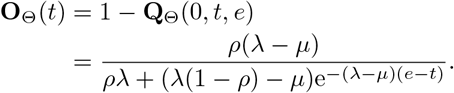

By construction, birth-death-sampling models are both Markovian and lineage-homogeneous.

## 3 Tree topologies

Tree topologies arising from diversification processes are binary (since a speciation event gives rise to a single new lineage under the models considered here) and rooted thus so are all the tree topologies considered here. Moreover, all the tree topologies considered below will be *labeled*, which means their tips, and consequently all their nodes, are unambiguously identified. From now on, “tree topology” has to be understood as “labeled-rooted-binary tree topology”.

Since the context will avoid any confusion, we still write 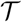 for the set of nodes of any tree topology 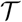. For all tree topologies 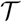, we put 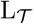 for the set of tips of 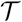. For all nodes *n* of 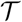, we note 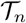 the subtree of 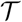 rooted at *n*.

For all sets *S*, |*S*| denotes the cardinality of *S*. In particular, 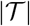 denotes the size of the tree topology 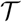 (i.e., its total number of nodes, internal or tips) and 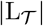 its number of tips.

### 3.1 Probability

Let us define 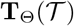 as the probability of a reconstructed tree topology 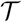 given its number of tips under a lineage-homogeneous diversification process.

#### Theorem 1 ([13]).

*Given its number of tips, the reconstructed tree topology* 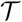 *of a realization of a lineage-homogeneous diversification process has probability* 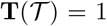 *if* 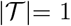, *i.e.*, 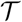 *is a single lineage. Otherwise, by putting a and b for the two direct descendants of the root of* 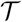, *the probability of the tree topology* 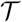 *is*

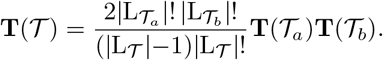

Assumptions of [13] are slightly different from those of Theorem 1 but its arguments still hold. The probability provided in [3, Supp. Mat., Appendix 2] is actually the same as that just above though it was derived in a different way from [13] and expressed in a slightly different form (see [4, Appendix 1]).

Theorem 1 implies in particular that 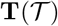 can be computed in linear time through a post-order traversal of the tree topology 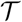.

### 3.2 Start-sets

A *start-set* of a tree topology 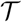 is a possibly empty subset *A* of internal nodes of 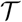 which is such that if an internal node of 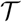 belongs to *A* then so do all its ancestors. Remark that, basically, the empty set ∅ is start-set of any tree topology and that if *A* and *A′* are two start-sets of 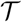 then both *A* ∪ *A′* and *A* ∩ *A′* are start-sets of 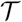.

Being given a tree topology 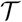 and a non-empty start-set *A*, we define the *start-tree* 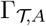 as the subtree topology of 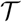 made of all nodes in *A* and their direct descendants. By convention, 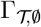, the start-tree associated to the empty start-set, is the subtree topology made only of the root of 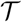.

For all tree topologies 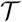, we define

- 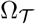 as the set of all start-sets of 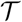, and for all internal nodes *n*,
- 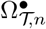 as the set of all start-sets *A* of 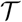 such that *n* ∈ *A*,
- 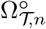 as the set of all start-sets *A* of 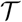 such that *n* ∉ *A*, and
- 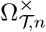 as the set of all start-sets *A* of 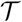 such that *n* is a tip of 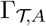.

Figure 2 displays examples of start-trees and of sets of start-sets of the types above.

**Figure 2:**
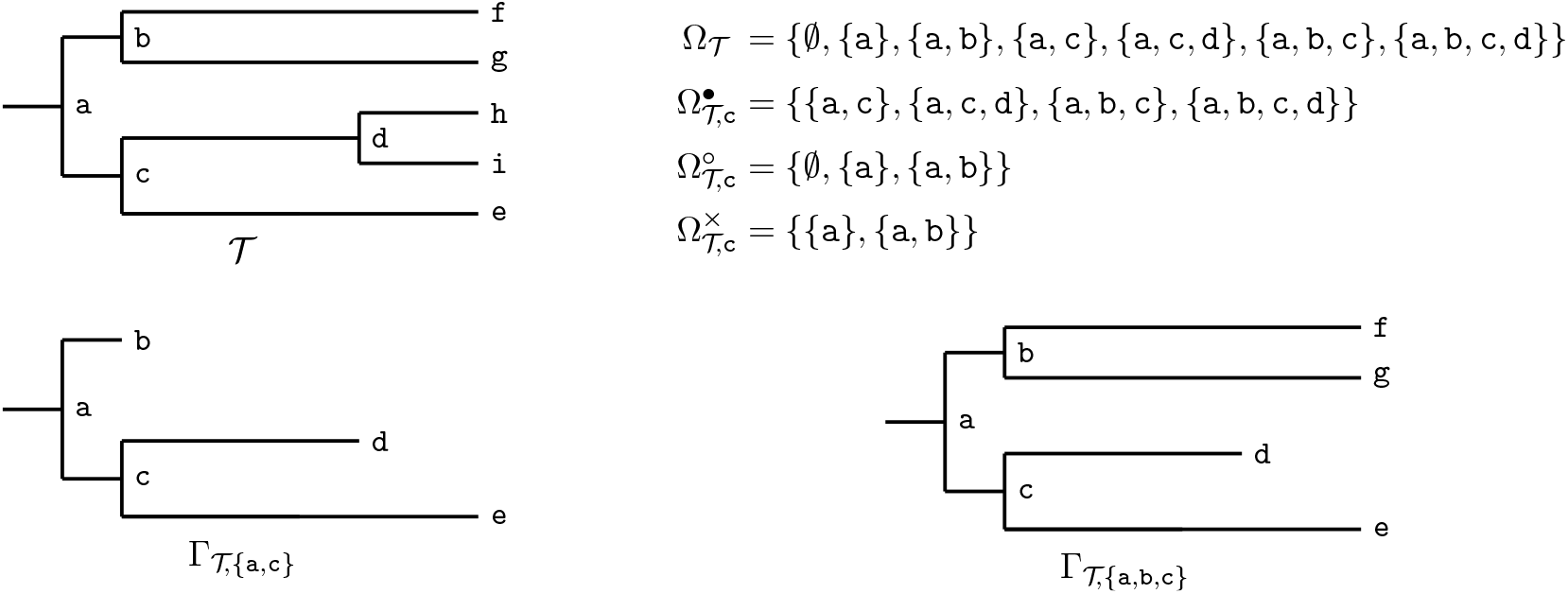
A tree 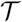 (top-left), examples of sets of start-sets of 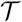 of various types (top-right) and the start-trees of 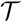 associated to the start-sets {`a`, `c`} and {`a`, `b`, `c`} (bottom).

## 4 Patterns

In this section, we shall consider diversification processes starting at origin time *s* and ending at time *e* following a birth-death-sampling model Θ = (*s, e, λ, μ, ρ*). A *pattern* is a part of the observed diversification process starting from a single lineage at a given time and ending with a certain number of lineages at another given time, these ending lineages being either observable or *special*, where “special” means “only known to be alive at the end time (and selected for some reason in the dataset)”. It consists of a 3-tuple 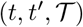 where *t* and *t*′ are the start and end times of the pattern and 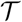 is the resulting tree topology. We shall consider two types of patterns: standard and special patterns. Standard patterns ends with only observable lineages. All the ending lineages of a special pattern are observable except one which is special (Fig. 3). Standard and special patterns are very similar to patterns defined in [3] for the fossilized-birth-death process.

**Figure 3:**
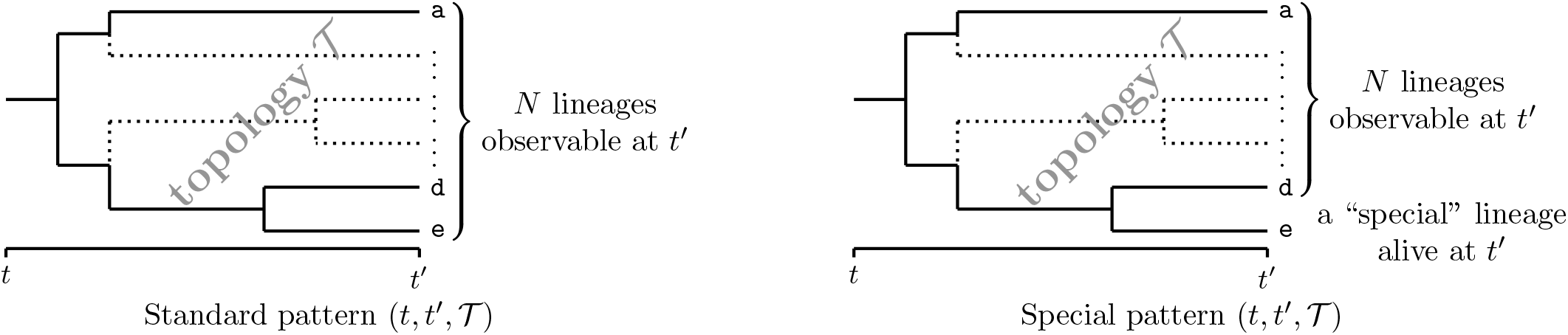
The two types of patterns used to compute probability distributions.

### 4.1 Standard patterns

#### Definition 1.

*A standard pattern* 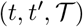 *starts with a single lineage at time t and ends with a tree topology* 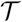 *and* 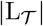 *observable lineages at time t′ (Fig. 3-left)*.

Let us compute the probability **X**_Θ_(*t, t′, N*) that a single lineage at time *t* ∈ [*s, e*) has *N* descendants observable at time *t′* ∈ (*t, e*] under the diversification model Θ. This probability is the sum over all numbers *j ≥* 0, of the probability that the lineage at *t* has *j* + *N* descendants at *t*′ in the whole process, which is equal to **Q**_Θ_(*t, t′, j* + *N*), among which exactly *N* ones are observable (i.e., 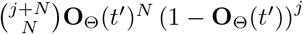). Under the diversification model Θ, we thus have

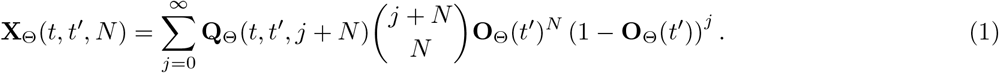

If Θ is the birth-death-sampling model (*s, e, λ, μ, ρ*), Equation 1 becomes

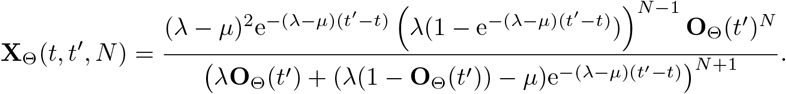

The probability of the standard pattern 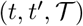 is the probability of the tree topology 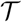 conditioned on its number of tips, which is 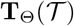 multiplied by the probability of observing this number of tips in a standard pattern, which is that of getting 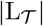 observable lineages at *t*′ from a single lineage at *t*, i.e., 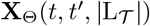.

#### Claim 1.

*Under the diversification model* Θ, *the probability of the standard pattern* 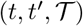 *with s ≤ t < t′ ≤ e is*

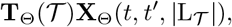

*where* **T**_Θ_ = **T** *if* Θ *is lineage-homogeneous*.

### 4.2 Special patterns

#### Definition 2.

*A special pattern* 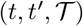 *starts with a single lineage at time t* ∈ [*s, e*) *and ends with the tree topology* 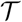 *at t′* ∈ (*t, e*], *thus with* 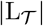 *descendants at t′ among which* 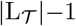 *are observable and one is a distinguished “special” lineage of fate a priori unknown after t′(Fig. 3-right).*

Let us now compute the probability **Y**_Θ_(*t, t′, N* + 1) that a single lineage at time *t* ∈ [*s, e*) has one special descendant and *N* descendants observable from *e* at time *t′* ∈ (*t, e*]. This probability is the sum over all numbers *j*, of the probability that the lineage at *t* has *j* + *N* + 1 descendants at *t*′ in the whole process, which is equal to **Q**_Θ_(*t, t′, j* +*N* +1), among which the special one is picked, exactly *n* ones are observable and *j* ones are not observable, which leads to 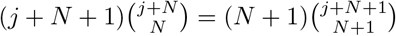 possibilities. Under the diversification model Θ, we have that

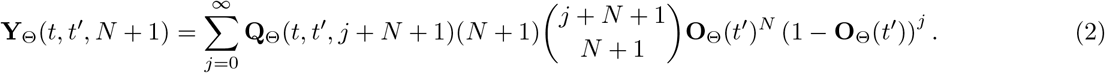

If Θ is the birth-death-sampling model (*s, e, λ, μ, ρ*), Equation 2 gives us that

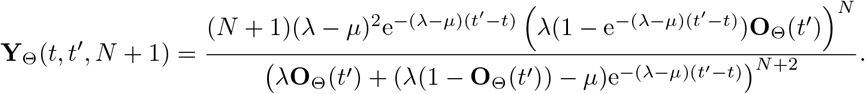

The probability of the special pattern 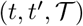 is the probability of the tree topology 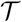 conditioned on its number of tips, which is 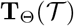 multiplied by the probability of observing this ending configuration in a special pattern, i.e., 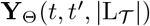.

#### Claim 2.

*Under the diversification model* Θ, *the probability of the special pattern* 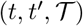 *with s ≤ t < t ′ ≤ e is*

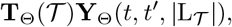

*where* **T**_Θ_ = **T** *if* Θ *is lineage-homogeneous.*

## 5 Probability densities of topologies with temporal constraints and shifts

The probability 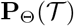 of observing a tree topology 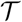 under a diversification model Θ with origin and end times *s* and *e* is that of the corresponding standard pattern, i.e., we have that

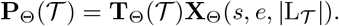

We shall see in this section how to compute the probability of a tree topology under the constraint that some of its divergence times are known to be anterior or posterior to given times.

### 5.1 Temporal constraints

Let us put *τ_n_* for the (random variable associated to the) divergence time corresponding to the node *n* of 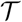. Being given internal nodes 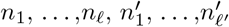 of 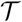 and times 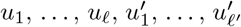 between *s* and *e* (both not included), we aim to compute the joint probability of 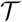 and of observing 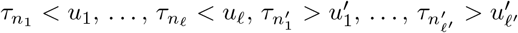, under the model Θ, i.e.,

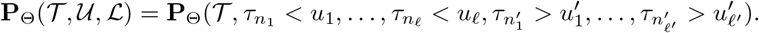

The temporal constraints induced by the tree topology, i.e., that we have necessarily *τ_n_ ≤ τ_m_* if *n* is an ancestor of *m* are implicitly assumed granted in the probability above. The constraints 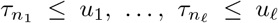 will be referred to as *upper temporal constraints* and summarized as the set of pairs “node-time” 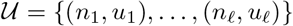, and the constraints 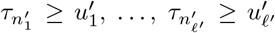, will be referred to as *lower temporal constraints* and summarized as the set of pairs 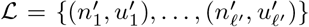. We assume that the temporal constraints are consistent with another (otherwise they would basically lead to a null probability). For all subsets of internal nodes 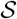 of 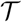, we write 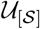 (resp. 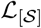) for the set of upper (resp. lower) temporal constraints of 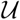 (resp. of 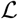) involving nodes in 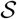, namely 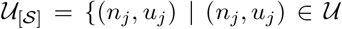 and 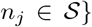 (resp. 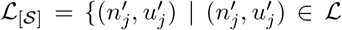 and 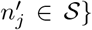). For all times *t*, we define 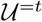 (resp. 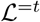) as the set of temporal constraints of 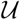 (resp. 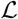) involving *t*, namely, 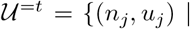 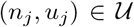 and *u*_*j*_ = *t*} (resp. 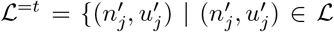 and 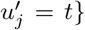). In the same way, we define 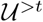 and 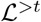 as the subsets of temporal constraints of 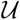 and 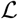 respectively, which involved times strictly posterior to *t*, i.e., 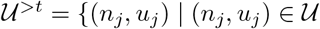 and *u_j_ > t*} and 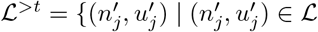 and 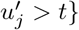.

#### Theorem 2.

*Let* 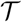 *be a tree topology,* Θ *be a Markovian diversification model from origin time s to end time e and* 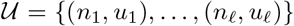 *and* 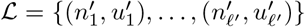 *be two sets of upper and lower temporal constraints respectively. Let o be the oldest time involved in a temporal constraint or the end time if there are none, namely*,

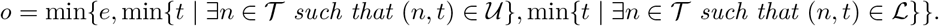

*Let us define the set* 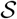 *of internal node subsets of* 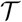 *as the intersection of*

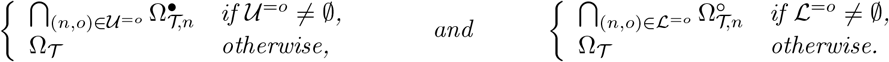

*The joint probability* 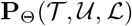 *of observing the tree topology 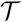 with the temporal constraints 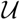 and 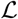 under* Θ *verifies*

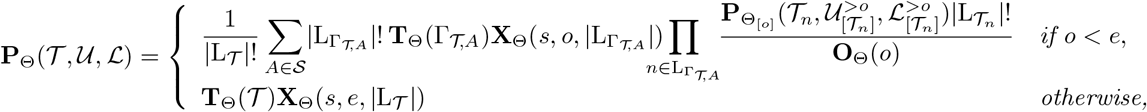

*where* Θ_[*o*]_ *is the model* Θ *restricted to the time interval* [*o, e*].

*Proof.* Appendix C.1.

The general idea of the computation presented in Theorem 2 is first to consider the oldest time involved in a temporal constraint (referred to as *o* in Theorem 2 and which is *t* in Figure 4), then to consider the part of the diversification process which occurred before the oldest time and the part(s) which occurred after the oldest time.

**Figure 4:**
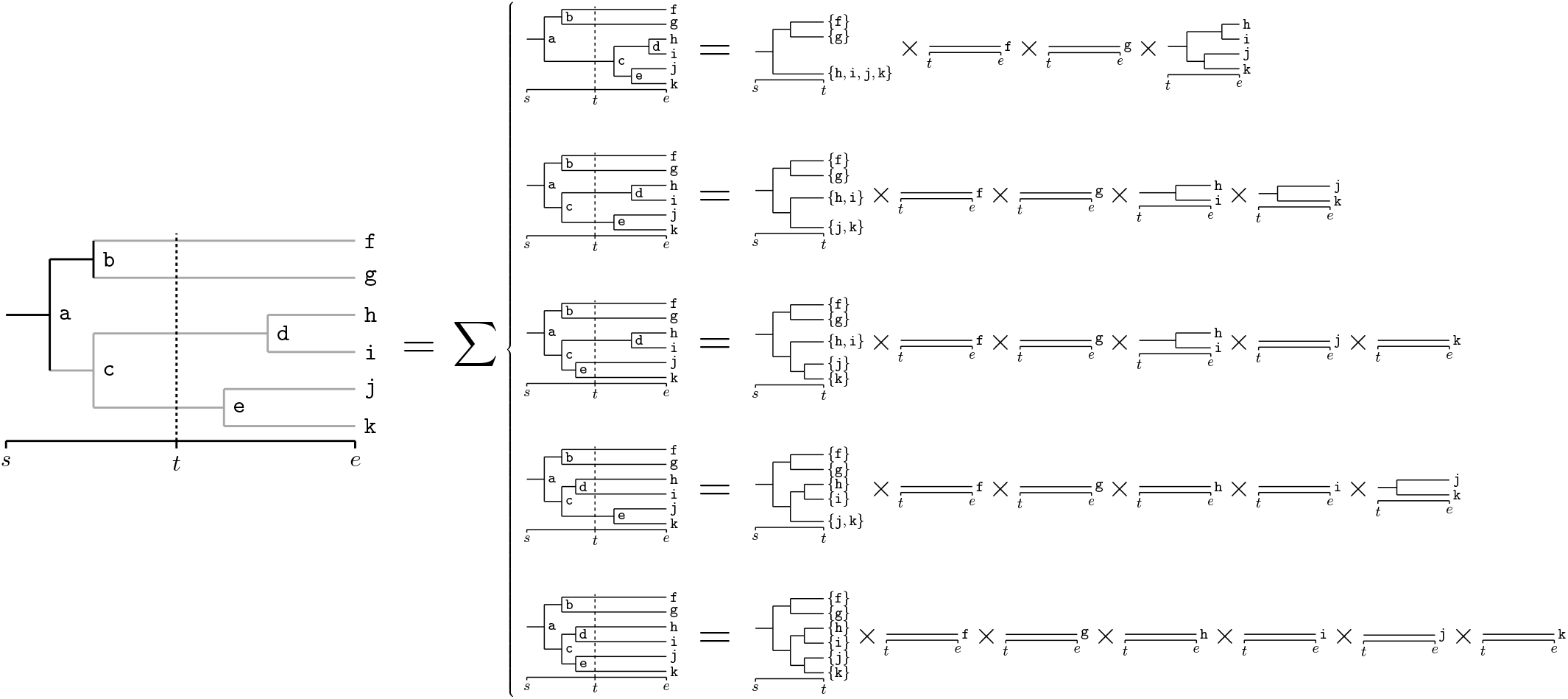
Schematic of the computation of the probability 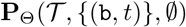, i.e., that the divergence time associated with node b is strictly anterior to *t*. Under the notations of Theorem 2, we have that 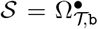. Nothing is known about divergence times in the gray part of the tree at the left. The only information about divergence times in black parts of all trees is their relative position with regard to *t*.

An issue here is that these parts are a priori unknown since, by construction, they are determined by the relative positions of the divergence times with regard to the oldest time. If some of the divergence times are directly or indirectly (from their ancestors or their descendants) constrained, thus known, to be posterior or anterior to the oldest time, some of them may be not and we have to consider all the possibilities consistent with the temporal constraints as displayed in the left-hand side of the second column of Figure 4. Remark that since all these possibilities are mutually exclusive, their respective probabilities can be summed in order to obtained the probability of the initial tree topology with the given temporal constraints.

In order to compute the probability of each possibility (i.e., in which all the relative positions of the divergence with regard to the oldest time are fixed), we use the Markov property to write the probability of the tree topology with the temporal constraints as the product of the probability of the part of the diversification which happened before the oldest time (which by construction contains no temporal constraint and therefore is a standard pattern) to the probabilities of the parts which happened after the oldest time which is a set of independent tree topologies starting from the oldest time with possibly some time constraints, which all involved times posterior to the oldest time. This part of the computation is illustrated by the five lines of Column 2 in Figure 4, in which the right-hand sides start with standard patterns corresponding to the diversification anterior to the oldest time which are followed by a certain number of tree topologies starting from the oldest time (in this case without time constraints since the example contains only one constraint) corresponding to the diversification posterior to the oldest time. For all the possibilities, Claim 1 allows us to directly compute the part of diversification anterior to the oldest time.

Theorem 2 states that 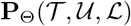 can be either calculated directly (if *o* = *e*) or expressed as a sum-product of probabilities of tree topologies with temporal constraints under birth-death-sampling models whose starting time is strictly posterior to the starting time of Θ, on which Theorem 2 can be applied and so on. Since each time that Theorem 2 is applied, we get tree topologies under models and temporal constraints in which the starting time has been discarded, we eventually end up in the case where the oldest time is the end time of the diversification for which the probability can be calculated directly. To summarize, the probability 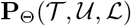 can be computed by recursively applying Theorem 2.

### 5.2 Shifts

We shall see how to compute the probability of a tree topology 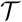 under a diversification model Θ starting at *s* and ending at *e* by assuming that one of its clades follows another diversification model 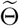 from a given time *t* ∈ [*s, e*] to the end time *e*. Note that this implicitly assumes that the lineage originating this particular clade was alive at *t* (Fig. 5).

**Figure 5:**
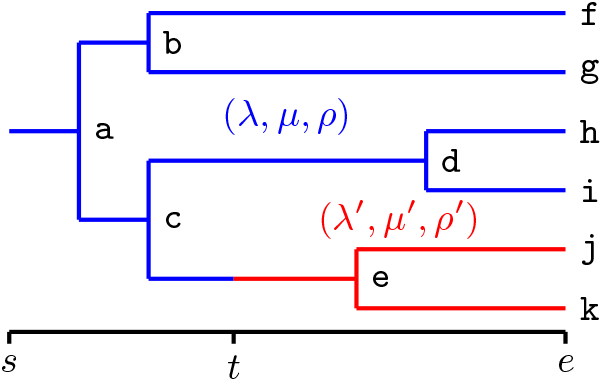
A tree topology with a shift at time *t* for the clade `{e, j, k}`

#### Theorem 3.

*Let* 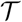 *be a tree topology, s ≤ t ≤ e be three times*, Θ *and* 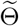 *be two Markovian diversification models from origin times s and t respectively and both to end time e, and m be a node of 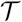. By putting* Θ_[*t*]_ *for the model* Θ *restricted to* [*t, e*], *the probability* 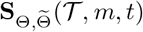 *of observing the tree topology* 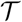 *assuming that evolution follows* Θ *on 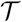 except on 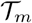 on which it follows* 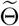 *from time t verifies*

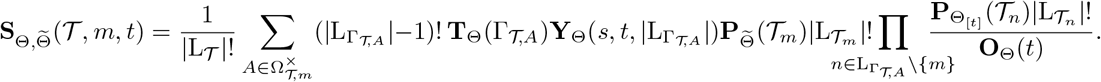

*Proof.* Appendix C.2.

The idea of the proof is essentially the same as that of Theorem 2.

Let us remark that the trees starting from *t* are standard patterns. It follows that 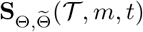 can be equivalently written as

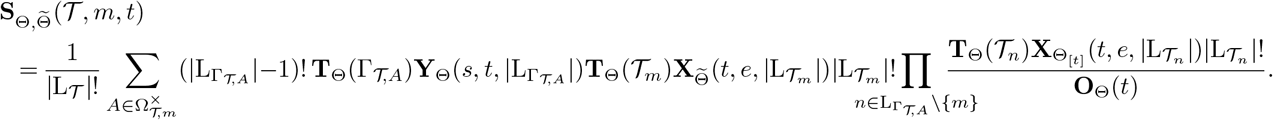

## 6 A quadratic computation

Since the number of start-sets may be exponential with the size of the tree, notably for balanced trees, Equations of Theorems 2 and 3 do not directly provide a polynomial algorithm for computing the probabilities considered in these theorems. In the case where the diversification is lineage-homogeneous, the form of the probability of the tree topology conditioned on its number of tips provided by Theorem 1 allows us to factorize the computation of Theorem 2 (and of Theorem 3) in order to obtain a polynomial algorithm. Let us sketch the general idea of this computation.

In the case where the diversification model Θ is lineage-homogeneous, Theorem 1 implies that **T**_Θ_ = **T**. Let us assume that the temporal constraints are such that 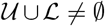 and let *a* and *b* be the two direct descendants of the root of 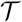. For all start-sets *A* of 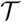, we define 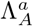 and 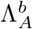 as the subtrees of 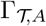 rooted at *a* and *b* respectively, namely 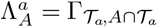 and 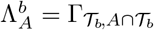. From Theorems 2 and 1, we have that

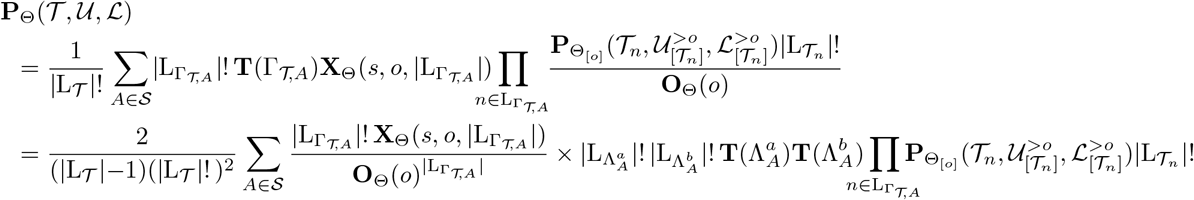

Since by construction a tip of 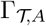 is either a tip of 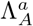 or a tip of 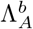, we have basically that

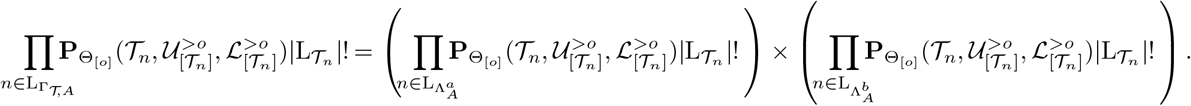

It follows that

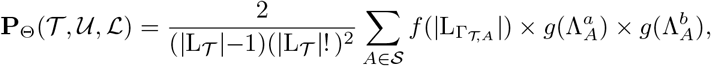

where

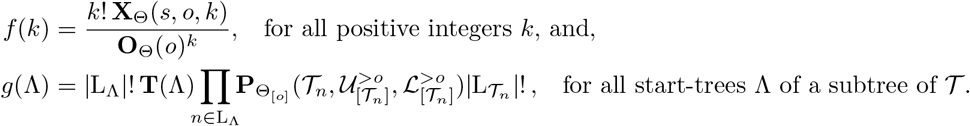

In plain English, computing 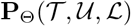 requires to sum over all start-sets *A* in 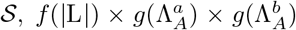, a product of three factors where the first one depends only on the number of tips of 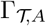 and the two following ones depend on the subtrees 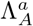 and 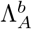 respectively (i.e., on the start-set *A* and on the subtrees 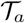 and 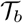).

Let us start by regrouping the terms of the sum with respect to the number of tips of the corresponding start-trees:

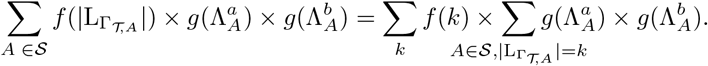

Next, we put 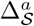 (resp. 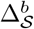) for the set of subtrees rooted at *a* (resp. at *b*) of the start-trees 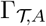 for all 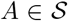, namely 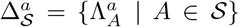 and 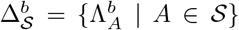. Moreover, for all 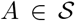 and since 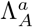 and 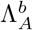 are the two subtrees pending from the root of 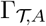, we have that 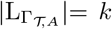 if and only if 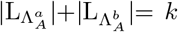. For all *k*, a set *A* belongs to 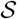 and is such that 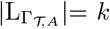 if and only if there exist a tree 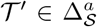 and a tree 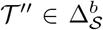 such that *A* is the union of the root of 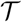 and of nodes of 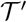 and 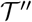 and 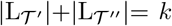. It follows that the terms of 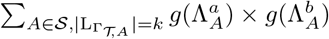 can be regrouped and factorized in order to get that

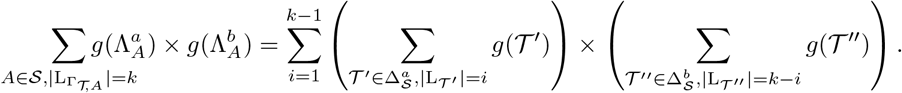

To summarize, if one assume that the quantities 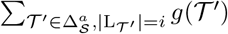 and 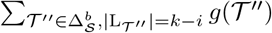 are known, we have written 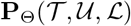 as a sum of a quadratic number of terms, namely

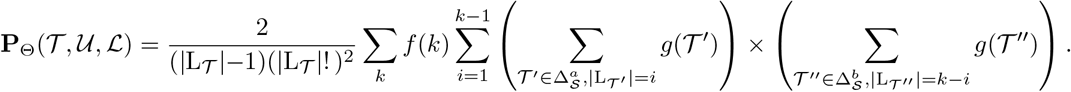

Appendix C.3 shows how the quantities 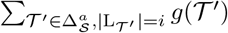 and 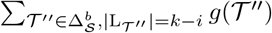, which are referred to as **W**_*a,i*_ and **W**_*b*,*k*−*i*_ respectively in the appendix, can be recursively computed in order to obtain a computation with total complexity quadratic with the size of 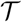. We eventually obtain the following theorem.

### Theorem 4.

*Let* 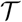 *be a tree topology*, Θ *be a lineage-homogeneous Markovian diversification model and* 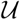 *and* 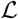 *be two sets of upper and lower temporal constraints respectively. If the probability of the ending configuration of any standard pattern can be computed in constant time then the probability* 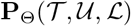 *can be computed with complexity O*(1) *both in time and memory space if* 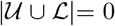 *(i.e., if there are no constraints) and with time complexity* 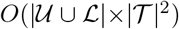 *and memory space complexity* 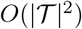 *otherwise*.

*Proof.* Appendix C.3

Theorem 4 holds in particular for birth-death-sampling models and sampled-generalized-birth-death models presented in Appendix B.

It can be proved in the same way that the shift probability 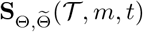 of Theorem 3 can be computed with time and memory space complexity 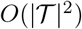 if the probability of the ending configuration of any standard or special pattern can be computed in constant time and if the diversification process is lineage-homogeneous.

## 7 Divergence time distributions

We shall apply Theorem 2 to compute divergence time distributions of tree topologies with temporal constraints.

### Claim 3.

*Let* 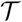 *be a tree topology,* Θ *be a diversification model from origin time s to end time e*, 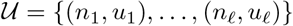 *and* 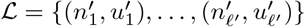 *be two sets of upper and lower temporal constraints respectively and m be an internal node of 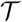. The probability that the divergence time τ_m_ associated with m is anterior to a time t* ∈ [*s, e*] *conditioned on observing the tree topology 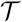 with the temporal constraints 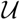 and 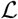 under* Θ *is*

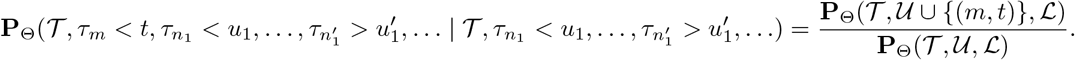

The computation of the divergence time distributions was performed on a contrived tree topology and on the Hominoidea subtree. Results are displayed in Figures 6 and 7 where the probability densities are computed from the corresponding distributions by finite difference approximations.

**Figure 6:**
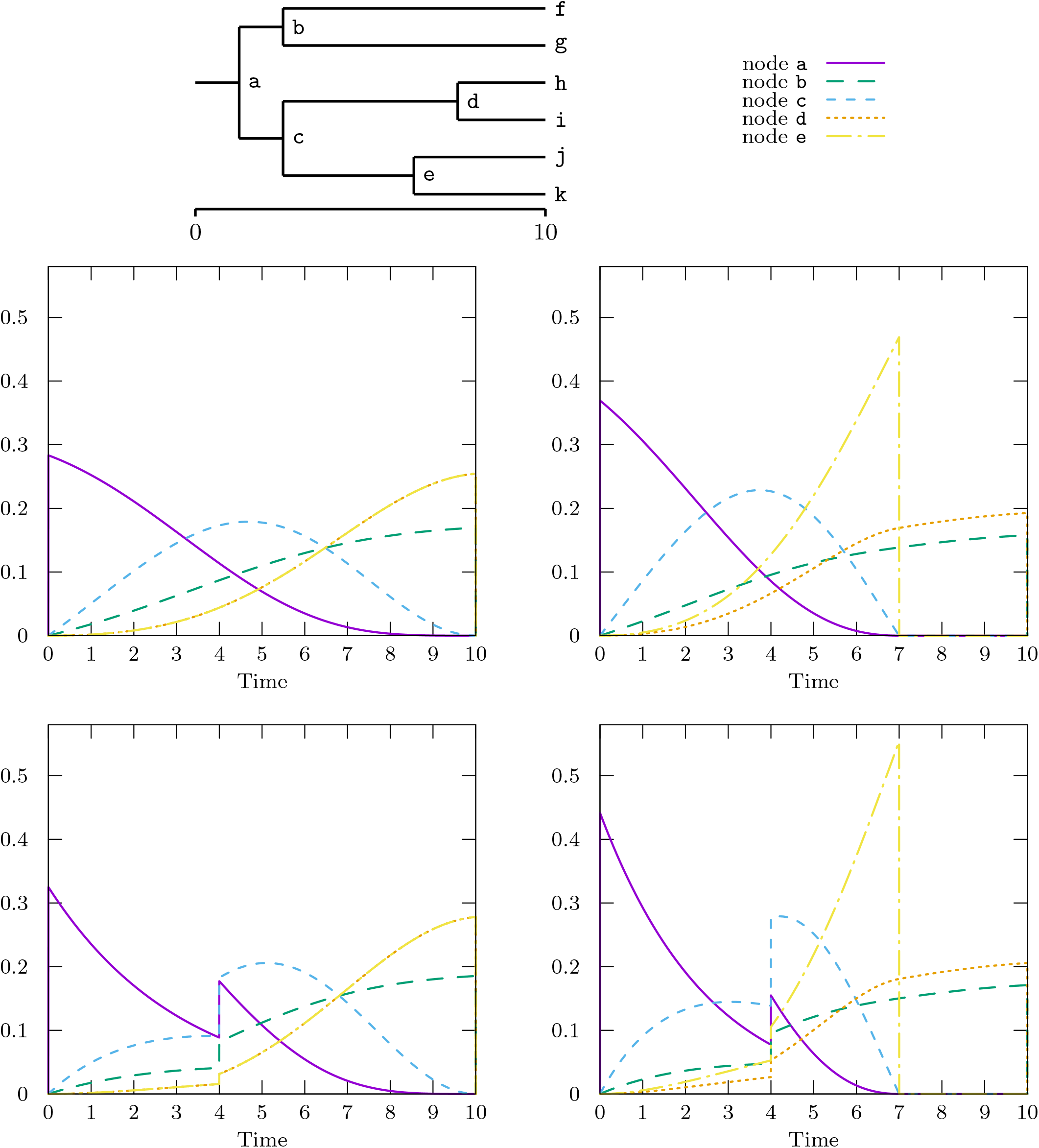
Divergence time probability densities of the tree displayed at the top, in the first row of plots by assuming a diversification process running from time 0 to 10 under a birth-death-sampling model with parameters *λ* = 0.2, *μ* = 0.02 and *ρ* = 0.5 between times 0 and 10 and in the second row of plots by assuming a piecewise constant birth-death-sampling model with parameters *λ*_0_ = 0.1, *μ*_0_ = 0.02 and *ρ*_0_ = 0.1 between times 0 and 4 (only 10% of the lineages survives to time 4) and parameters *λ*_1_ = 0.2, *μ*_1_ = 0.02 and *ρ*_1_ = 0.5 between times 4 and 10. Plots of the first column are computed with no constraint on the divergence times and those of the second column by constraining the divergence time associated to node `e` to be anterior to 7. Densities of nodes `d` and `e` are confounded in the plots of the first column. Densities was obtained from the corresponding distributions by finite difference approximations.

**Figure 7:**
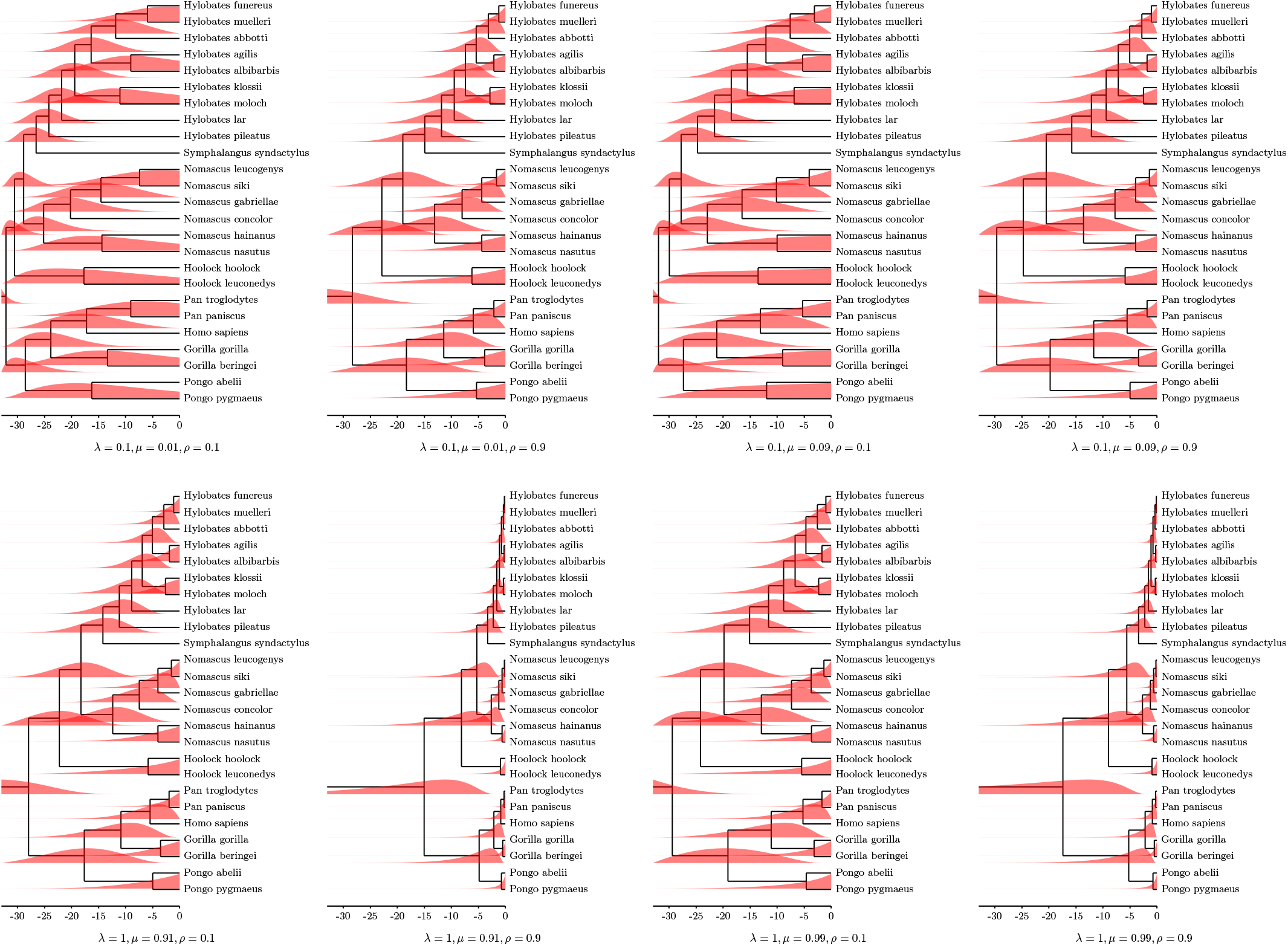
Divergence time probability densities of the Hominoidea tree from [7] under birth-death-sampling models with parameters *λ* = 0.1 or 1, *μ* = 0.01 or 0.09 and *ρ* = 0.1 or 0.9. Internal nodes are positioned at their median divergence time. Densities was obtained from the corresponding distributions by finite difference approximations.

Figure 6 shows how considering models which are not time-homogeneous such as the piecewise-constant-birth-death models and adding temporal constraints on some of the divergence times influences the shapes of the divergence times distributions of all the nodes of the tree topology. In particular, divergence time distributions may become multimodal, thus hard to sample. Let us remark that a temporal constraint on the divergence time of a node influences the divergence time distributions of the other nodes of the tree topology, even if they are not among its ascendants or descendants.

In order to illustrate the computation of the divergence time distributions on a real topology, let us consider the Hominoidea subtree from the Primates tree of [7]. The approach can actually compute the divergence time distributions of the whole Primates tree of [7] but they cannot be displayed legibly because of its size.

The divergence time distributions were computed under several (simple) birth-death-sampling models, namely all parameter combinations with *λ* = 0.1 or 1, *μ* = *λ* − 0.09 or *λ* − 0.01 and *ρ* = 0.1 or 0.9. Since the difference *λ* − *μ* appears in the probability formulas, several sets of parameters are chosen in such a way that they have the same difference between their birth and death rates.

Divergence time distributions obtained in this way are displayed in Figure 7 around their internal nodes (literally, since nodes are positioned at the median of their divergence times). Each distribution is plotted at its own scale in order to be optimally displayed. This representation allows us to visualize the effects of each parameter on the shape and the position of distributions, to investigate which parameter values are consistent with a given evolutionary assumption etc.

Birth-death-sampling models are not identifiable, since several sets of parameters leads to the same probability distributions. Namely, *ρ* and *ρ′* being two sampling probabilities, if one sets 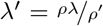 and 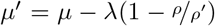, the probability densities of any phylogenetic tree (by considering or not considering its divergence times) is the same under the model (*s*, *e*, *λ*, *μ*, *ρ*) as under the model (*s*, *e*, *λ*′, *μ*′, *ρ*′) [31]. An identifiable parametrization of birth-death-sampling models is provided in [31].

We observe on Figure 7 that, all other parameters being fixed, the greater the speciation/birth rate *λ* (resp. the sampling probability *ρ*), the closer are the divergence time distributions to the end time.

Influence of the extinction/death rate on the divergence time distributions is more subtle and ambiguous, at least for this set of parameters. All other parameters being fixed, it seems that an increase of the extinction rate tends to push distributions of nodes close to the root towards the starting time and, conversely, those of nodes close to the tips towards the end time.

The divergence time distributions obtained for *λ* = 0.1, *μ* = 0.01 and *ρ* = 0.9 (Fig. 7, column 2, top) and for *λ′* = 1, *μ′* = 0.91 and *ρ′* = 0.1 (Fig. 7, column 1, bottom) are very close one to another. The same remark holds for *λ* = 0.1, *μ* = 0.09 and *ρ* = 0.9 (Fig. 7, column 4, top) and for *λ′* = 1, *μ′* = 0.99 and *ρ′* = 0.1 (Fig. 7, column 3, bottom). This certainly comes from identifiability issue of the birth-death sampling model since in both cases we have that 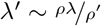 and 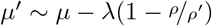.

The variety of shapes of divergence times probability densities observed in Figures 6 and 7 exceeds that of standard prior distributions used in phylogenetic inference, e.g., uniform, lognormal, gamma, exponential [16, 14].

### 7.1 A previous approach

A previous approach for computing the probability density of a given divergence time is provided in [10]. It is based on the explicit computation of the probability density 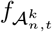 of the *k^th^* divergence time of a tree topology with *n* tips starting at *t* from the present, provided in [10], and the computation of the probability ℙ(*r*(*v*) = *k*) for the rank *r*(*v*) of the divergence time associated to the vertex *v* to be the *k^th^* which was given in [11]. The probability density *f_v_* of the divergence time associated to a vertex *v* of a tree topology with *n* tips is then given for all times *s* by

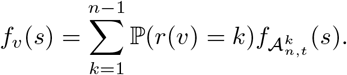

The probability density 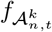 is computed in constant time and the probabilities ℙ(*r*(*v*) = *k*) for all nodes *v* are computed in a time quadratic with the size of the tree.

The computation of the probability density of the *k^th^* divergence time of tree relies on the fact that, under some homogeneity assumption, the divergence times are independent and identically distributed (iid) random variables. Approach provided in [10] was described in the case of birth-death models. It can be easily adapted to deal with piecewise-constant-birth-death-sampling models but extending this approach in order to compute divergence times distribution with temporal constraints requires to consider all the orders consistent with the set of constraints, which is certainly feasible but seems not straightforward.

### 7.2 An alternative method

By reviewing the present work, Amaury Lambert proposed an alternative method, which requires the property that the divergence times are iid random variables, to compute the probability that the divergence times of a tree topology satisfy a given set of temporal constraints. Although a presentation of this method is provided in his first review, let us sketch its idea.

We assume here that the diversification model starts from *s*, ends at *e* and is such that the divergence times are iid random variables which follow a distribution *F* which is known explicitly. This property is in particular granted for generalized-birth-death models [19] and for the model proposed in Appendix B. Let 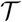 be a tree, 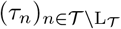 be the random variables associated to its divergence times and 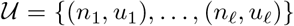 and 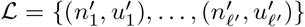 be a set of upper and lower time constraints respectively.

Amaury Lambert first remarks that the probability that 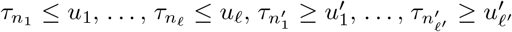 is equal to the probability that 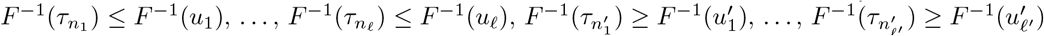, where the random variables 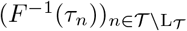 are independent and uniformly distributed over [0, 1].

Let us set *H_n_* = *F*^−1^(*τ*_*n*_) for all internal nodes *n* of 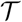 and

- 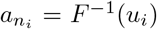 for all 1 ≤ *i* ≤ *ℓ* (i.e., for all nodes *n_i_* with upper time constraints in 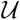) and *a_m_* = *F*^−1^(*e*) = 1 for all internal nodes *m* ∉ {*n*_1_,…, *n_ℓ_*},
- 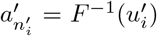 for all 1 ≤ *i* ≤ *ℓ*′ (i.e., for all nodes 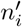 with lower time constraints in 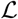) and 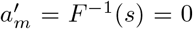 for all internal nodes 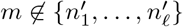.

Let us recall that the random variables 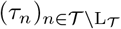, thus the uniform random variables 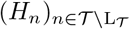, have to satisfy not only the constraints induced by 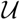 and 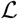 but also (implicitly) those deriving from the diversification process. In order to compute the probability that the independent and uniform random variables 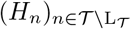 satisfy both the constraints induced by 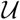 and 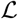 and those deriving from the tree topology, Amaury Lambert defines *Q_m_*(*x*) as the probability that the random variables 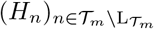 (i.e., those associated to the subtree 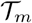) satisfy both these constraints and that *H_m_* ≥ *x*, for all nodes *m* of 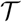 and all *x* ∈ [0, 1]. By setting *Q_n_*(*x*) = 1 for all tips *n* of 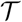 and all *x* ∈ [0, 1], the probability *Q_m_*(*x*) can be recursively computed for all internal nodes *m* from its two direct descendants *m*_1_ and *m*_2_, since from the independence property, we have that

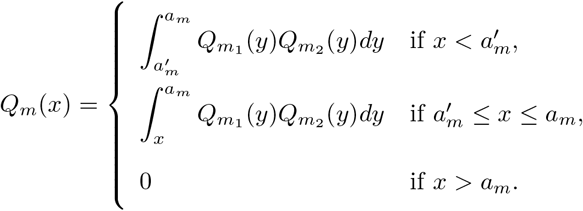

The probability that the random variables 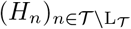 satisfy both the constraints induced by 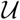 and 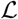 and those deriving from the tree topology, which is equal to the probability that the divergence times of 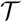 satisfy the temporal constraints 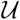 and 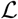, is finally given by 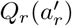, where *r* is the root of 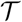.

By construction, the function *Q_m_*(*x*) is piecewise polynomial of degree smaller than 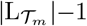 and its definition requires to consider a number of intervals bounded by the cardinality of the set of times involved in the temporal constraints of 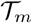. It follows that the symbolic computation of *Q_r_*(*x*) can be performed in 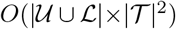, thus with exact same complexity as that of the algorithm presented in Section 6, though it certainly involves a smaller constant factor.

## 8 Direct sampling of divergence times

Theorems 2 and 4 and Claim 3 show how to compute the marginal (with regard to the other divergence times) of the divergence time distribution of any internal node of a phylogenetic tree from a given birth-death-sampling model. It allows in particular to sample any divergence time of the phylogenetic tree disregarding the other divergence times. We shall see in this section how to draw a sample of all the divergence times of any tree topology from a given birth-death-sampling model.

### Lemma 1.

*Let* 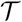 *be a tree topology of root r,* Θ *be a Markovian diversification model from origin time s to end time e. The probability that the root divergence time τ_r_ is anterior to a time t* ∈ [*s, e*] *conditioned on observing the tree topology* 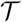 *under* Θ *is*

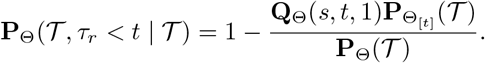

*Proof.* The probability that the divergence time *τ_r_* associated with *r* is anterior to a time *t* ∈ [*s, e*] is the complementary probability that *τ_r_* > *t*. Observing *τ_r_* > *t* means that the starting lineage at *s* has a single descendant observable at *t* from which descends the tree topology 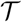 sampled at *e*. It follows that

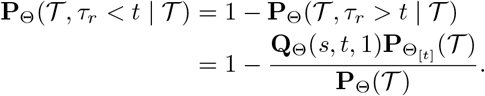

The probability 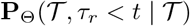 can be directly written as 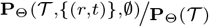. Lemma 1 shows that considering a temporal constraint is not necessary, which is particularly interesting in the birth-death-sampling case.

### Remark 1.

*Under the birth-death-sampling model* Θ = (*s*, *e*, *λ*, *μ*, *ρ*), *we have that*

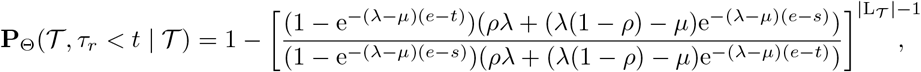

*which can be computed in constant time*.

Let us first show how to sample the divergence time of the root of a tree topology. The marginal, with regard to the other divergence times, of the distribution of the root-divergence time conditioned on the tree topology 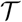 is the cumulative distribution function (CDF) 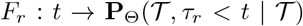. In order to sample *τ_r_* under this distribution, we shall use *inverse transform sampling* which is based on the fact that if a random variable *U* is uniform over [0, 1] then 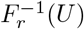 has distribution function *F_r_* (e.g., [2, chapter 2]). Since finding an explicit formula for 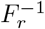 is not straightforward, we have to rely on numerical inversion at a given precision level in order to get a sample of the distribution *F_r_* from an uniform sample on [0, 1]. The current implementation uses the *bisection method*, which computes an approximate inverse with a number of *F_r_*-computations smaller than minus the logarithm of the required precision [2, p 32].

In order to sample the other divergence times, let us remark that by putting *a* and *b* for the two direct descendants of the root of 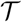 and *t* for the time sampled for the root-divergence, we have two independent diversification processes both starting at *t* and giving the two subtree topologies 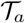 and 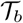 at *e*. By applying Lemma 1 to 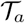 and 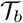 between *t* and *e*, the divergence times of the roots of these subtrees, i.e., *a* and *b*, can thus be sampled in the same way as above. The very same steps can then be performed recursively in order to sample all the divergence times of 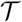. The time complexity of each sampling of a divergence time of 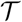 is obtained by multiplying the complexity of computing the probability of Lemma 1 with minus the logarithm of the precision required for the samples. From Remark 1, under the birth-death-sampling model Θ = (*s*, *e*, *λ*, *μ*, *ρ*), the computation of 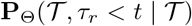 requires only the number of tips of 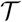 (in particular, the shape of 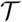 does not matter). In this case, the CDF *F_r_* can be computed at any time *t* with complexity *O*(1) and, with a pre-order traversal of 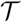, all its divergence times can be sampled in a time linear in 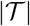 with a multiplicative factor proportional to minus the logarithm of the precision required for the samples.

The same approach can be applied in order to sample divergence times with temporal constraints and/or shifts.

## 9 Testing diversification shifts

Theorem 3 yields the computation of the probability density of a tree topology in which a given clade diversifies from a given “shift time” according a (simple) birth-death-sampling model different from that of the rest of the topology. This allows us to estimate the likelihood-ratio test for comparing the null model assuming a unique diversification model for the whole topology with the alternative model including a shift as displayed in Figure 5. Since the alternative model requires the implicit assumption that the lineage originating the “shifted” clade was alive at the shift time, we make the same assumption for the null model, i.e., the divergence time associated to the crown-node of the clade (resp. to the direct ancestor of the crown node) is assumed to be posterior (resp. anterior) to the shift-time. Basically, being given a tree topology, one of its clade and the shift time, we compute the ratio Λ_*N*_ of the maximum likelihoods of this topology with to without shift at the clade and shift time from Theorems 3 and 2 by using numerical optimization whenever a direct determination is not possible. Namely, in order to test a diversification shift at time *t* on the clade originating at node *m* of the tree topology 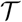, we consider the ratio

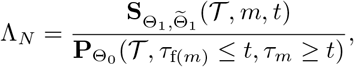

where f(*m*) is the direct ancestor of *m*, Θ_0_, Θ_1_, 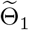 are diversification models with

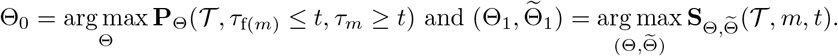

In order to assess the accuracy of Λ_*N*_, we compare it to three sister-group diversity tests considered in [36]. Namely, for two sister groups originating at shift time *t* with *N*_1_ > *N*_2_ terminal taxa and total sums of branch lengths *B*_1_ and *B*_2_ respectively, we have that

- the probability of observing this or greater difference between sister group diversities from [30] is 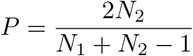,
- the likelihood ratio alternative provided in [28] is 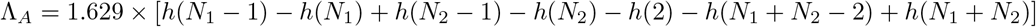, where 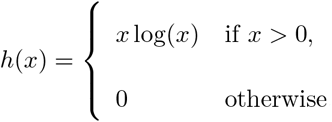
- the likelihood ratio from perfect-information given in [36] is 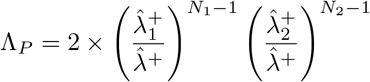, where 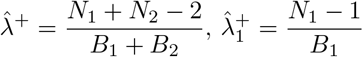 and 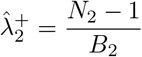.

I simulated topologies with and without shift according to pure-birth models, a.k.a. Yule models which are special cases of birth-death-sampling models with null death rate and full sampling, in the following way. Being given a general birth rate, a shift birth rate and the shift time, I first simulated topologies without shift from the general birth rate. Next, I filtered the simulated topologies by discarding those with less than 10 or more than 50000 nodes and those with a single lineage alive at the shift time. For each remaining simulation, I randomly picked a lineage alive at the shift time and replaced the clade originating from this lineage with a clade simulated with the shift rate from the shift to the end times in order to eventually obtain a topology with shift.

The quantities Λ_*N*_, the likelihood ratio obtained from Theorem 3, *P*, Λ_*A*_ and Λ_*P*_ are then evaluated with regard to their ability to discriminate between tree topologies with or without shift. Figure 8 displays the Receiver Operating Characteristic (ROC) plots obtained for all these quantities. We first observe that Λ_*N*_ significantly outperforms measures *P* and Λ_*A*_. In particular, in the case where the difference between the general and the shift birth rates is small (e.g., 0.6 and 1.0 in Fig. 8-left), performances of *P* and Λ_*A*_ are close to that of a random guess while Λ_*N*_ is still accurate. This was expected to at least some extent since Λ_*N*_ takes into account both the shift time and the whole tree topology while *P* and Λ_*A*_ are computed from the clade with the shift and its sister group. More surprisingly, Λ_*N*_ is only partially outperformed by Λ_*P*_, which is obtained from all the divergence times and the shift time. In the case where the general birth rate is 0.4 and the the shifted one is 1, the ability to distinguishes between phylogenies with or without diversification shift is almost as good with our likelihood ratio as with that of the perfect information. In the case where the general birth rate is 0.6, the likelihood ratio test Λ_*N*_ obtained from Theorem 3 outperforms the other tests for all positive discovery rates lower than 40%.

**Figure 8:**
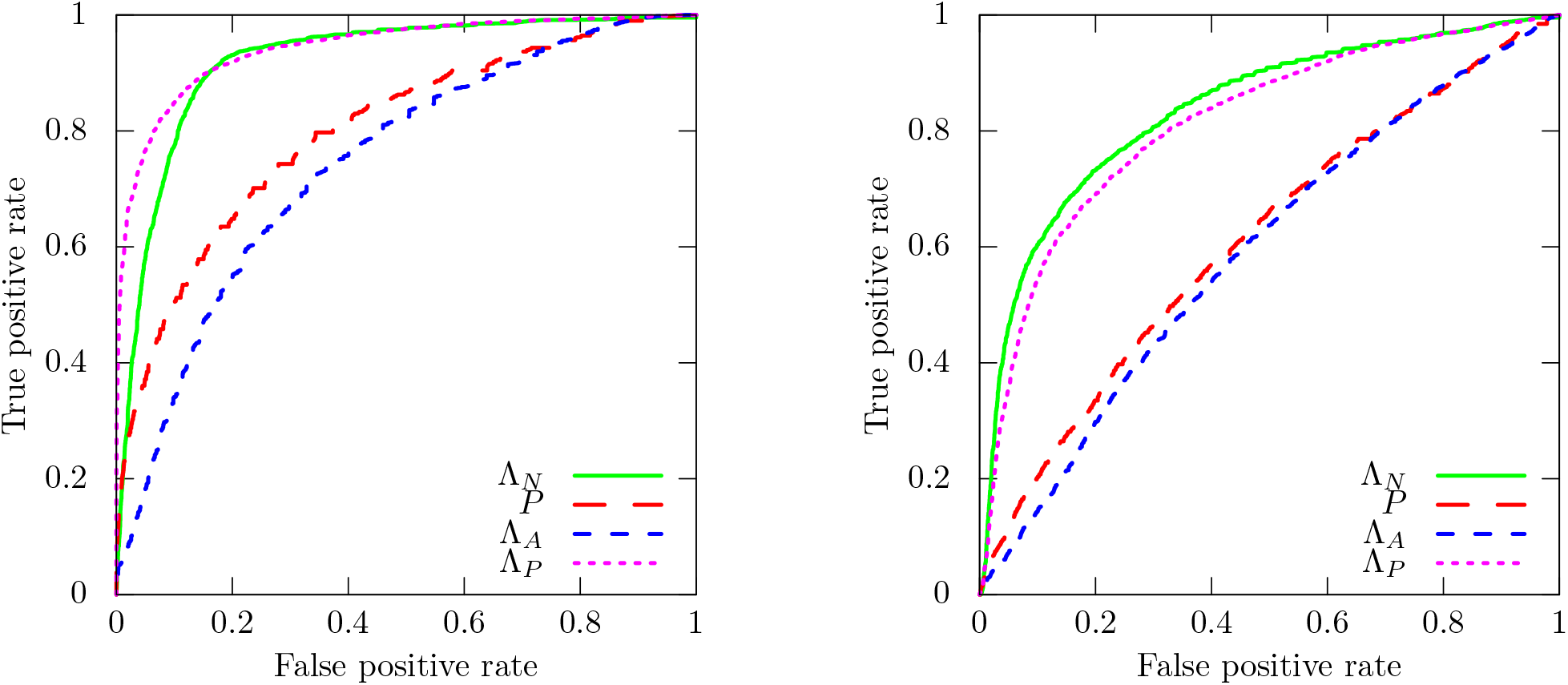
ROC plots of different measures for shift detection at left (resp. at right) are obtained by simulated 10000 Yule topologies with birth rate 0.4 (resp. 0.6) from times 0 to 10 and birth rate 1.0 from the shift time 5 to 10 for one of the clades present at time 5.

In order to illustrate the diversification tests on a biological dataset, let us consider the calibrated phylogeny of Cetacea from [29], which displayed in Figure 9. In [29], authors detected a diversification rate increase in Delphinidae using MEDUSA, a detection method developed in [1]. The general idea of MEDUSA is to fit birth and death models with increasing numbers of diversification shifts by stopping when the improvement in the Akaike Information Criterion (AIC) is smaller than a fixed threshold. Note that the MEDUSA method requires all the divergence times in order to fit the models [1].

**Figure 9:**
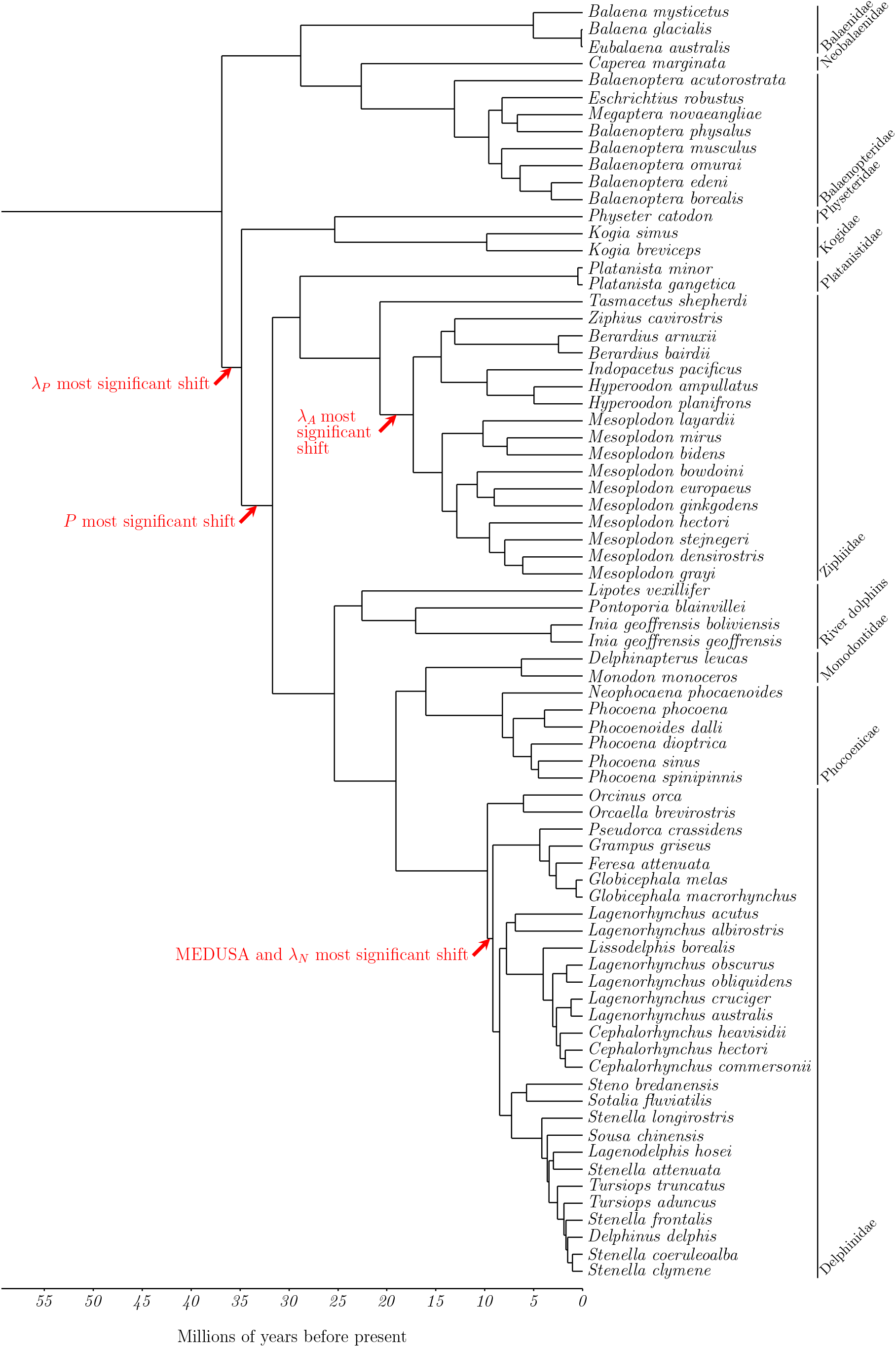
Time calibrated phylogeny of Cetacea [29].

We computed the quantities Λ_*N*_, *P*, Λ_*A*_ and Λ_*P*_ for all clades of the phylogeny of Cetacea, each time by setting the shift time to the time corresponding to the middle of the branch supporting the clade. Figure 9 displays the phylogenetic positions of the maxima observed for all these quantities. The maximal/most significant with regard to the likelihood ratio Λ_*N*_ was achieved at the position where the diversification rate increase was detected by MEDUSA [29, Fig. 1]. None of the other quantities *P*, Λ_*A*_ and Λ_*P*_ were maximal for this branch (Fig. 9).

## Acknowledgements

Version 4 of this preprint has been peer-reviewed and recommended by Peer Community In Evolutionary Biology (https://doi.org/10.24072/pci.evolbiol.100088). I warmly thank Amaury Lambert, Nicolas Lartillot, Dominik Schrempf and an anonymous reviewer for their useful comments and suggestions.

## A Table of the notations

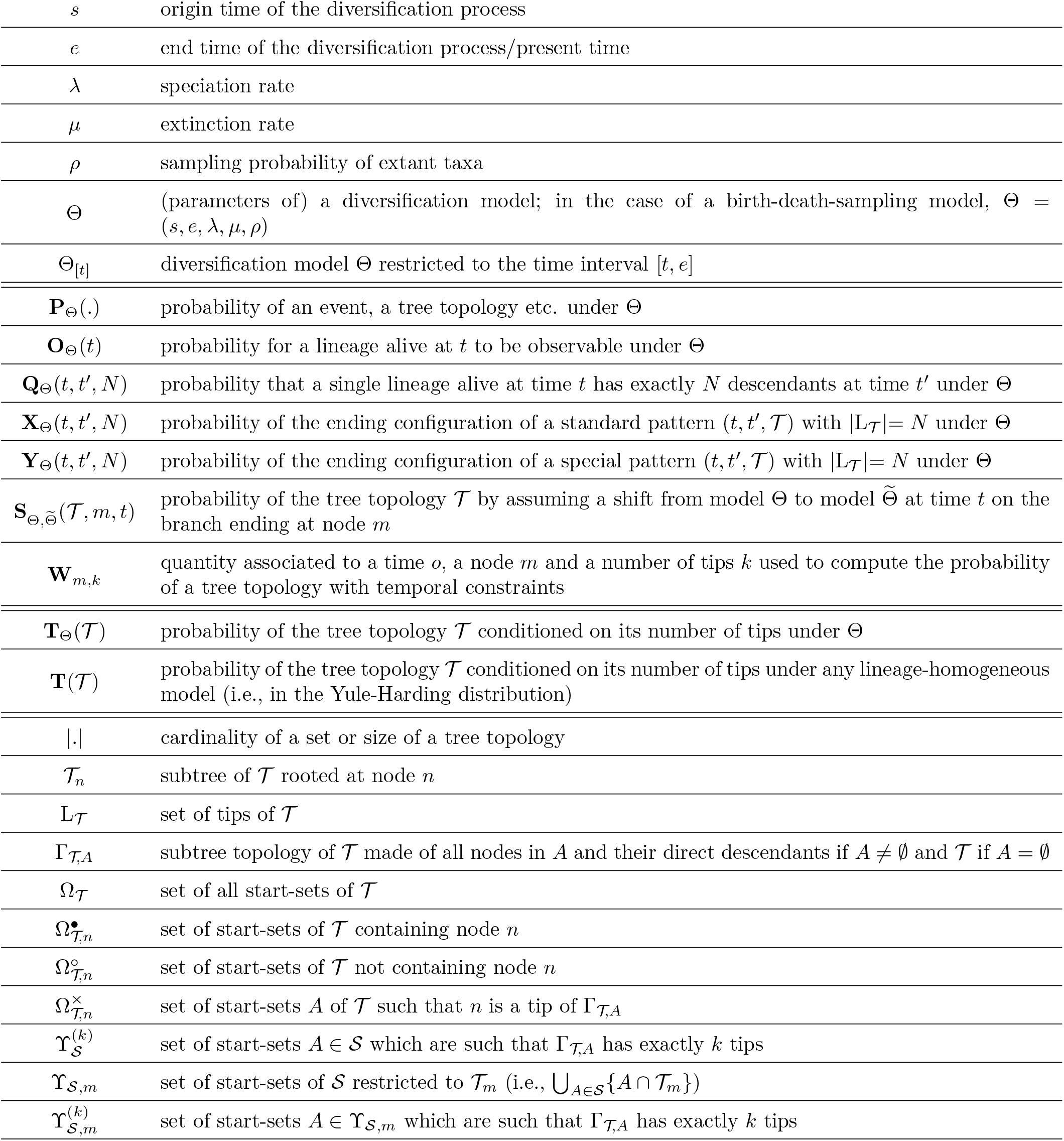

## B The generalized birth-death model with mass extinction events and extant sampling

The generalized birth-death process was introduced and studied in [17]. In this model, the speciation and extinction rates are allowed to change through time and are therefore given as two functions of the time, *λ*: *t* → *λ*(*t*) and *μ*: *t* → *μ*(*t*) (in this section, *λ* and *μ* denotes two functions of the time and are not real numbers like in Section 2.1). The probability 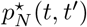 that a single lineage at time *t* has exactly *N* descendants at time *t′* by following the generalized birth-death (*λ, μ*) was given in [17]. We have that

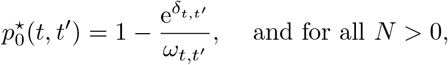

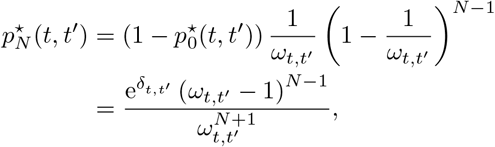

where 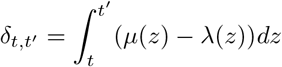 and 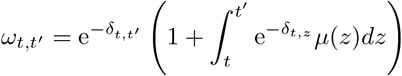.

**Figure 10:**
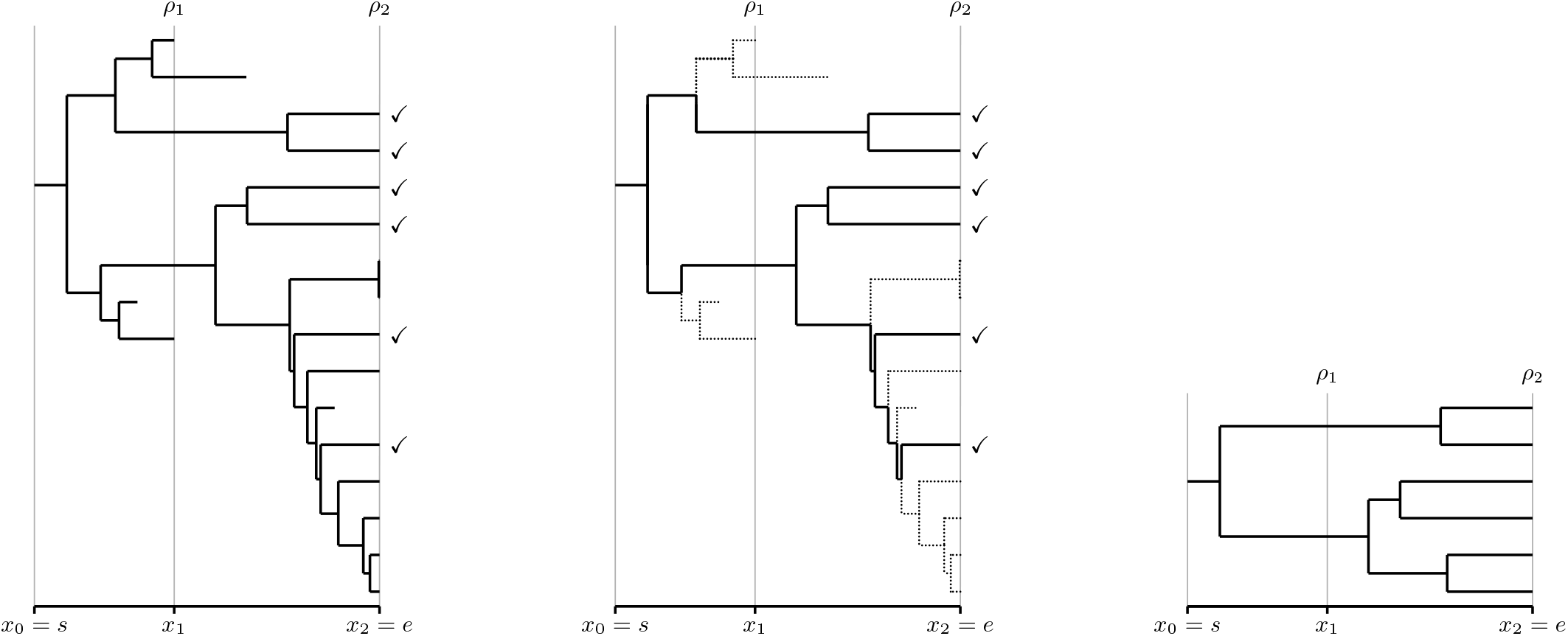
Left: the whole diversification process under the model (*s*, *e*, *λ*, *μ*, (*x*_1_, *ρ*_1_), (*x*_2_, *ρ*_2_)) (sampled extant species are those with ‘✓’); Center: the part of the process that can be reconstructed is represented in plain – the dotted parts are lost; Right: the resulting phylogenetic tree.

Following the idea of [32], we shall consider a more general model by allowing to uniformly sample lineages at a given set of times *x*_1_,…, *x_k_* with respective probabilities *ρ*_1_,…, *ρ_k_*. Namely, under the *sampled-generalized-birth-death* model Θ = (*s, e, λ, μ*, (*x_i_*, *ρ_i_*)_1≤*i*≤*k*_), lineages evolve following the generalized-birth-death model (*λ, μ*) between *s* and *e*, the origin and end times of the diversification process, and are uniformly sampled with probability *ρ_i_* at each time *x_i_* for 1 ≤ *i* ≤ *k* (Fig 10). In practice, sampling lineages at a time *x_i_* anterior to the present time has to be interpreted as a mass extinction event (a lineage not sampled at *x_i_* is assumed to have become extinct exactly at *x_i_*) while sampling at the present time accounts for our incomplete knowledge of extant species (a species not sampled at the present time is assumed unknown). From now on, we assume without loss of generality that the last sampling time is the end/present time, i.e., *x_k_* = *e*, and we set *x*_0_ = *s*, the origin time of the diversification process. Like in Section 2.1, we are interested in the reconstructed process, i.e., the part of the process which is observable from the present/end time (Fig 10).

By construction, sampled-generalized-birth-death models are both Markovian and lineage-homogeneous. Extending the approaches which are presented in the main text in order to deal with the sampled-generalized-birth-death model Θ = (*s, e, λ, μ*, (*x_i_, ρ_i_*)_1≤*i*≤*k*_) only requires to compute the probabilities of the ending probabilities of standard and special patterns, which can be obtained from the probabilities **O**_Θ_(*t*) and **Q**_Θ_(*t, t′, N*) for all positive numbers *N* and all times *s* ≤ *t* ≤ *t′* ≤ *e*. Let us see how to compute these last two probabilities.

In order to avoid ambiguity, we put *t*^+^ (resp. *t*^−^) for “an infinitesimal time after (resp. before) the time *t*”. In particular, time 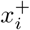 (resp. 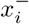) is immediately after (resp. before) the *i*^th^ sampling. By convention, we set **Q**_Θ_(*t*, *t′*, *N*) = **Q**_Θ_(*t*, (*t′*)^+^, *N*), i.e., **Q**_Θ_(*t*, *t′*, *N*) is the probability that a single lineage at time *t* has *N* descendants immediately after *t′*.

In order to compute **Q**_Θ_(*t, e*, 0), the probability that a lineage alive at *t* has no sampled descendant at the end/present time *e* = *x_k_* under the sampled-generalized-birth-death model Θ = (*s, e, λ, μ*, (*x_i_, ρ_i_*)_1≤*i*≤*k*_), let us remark that for all 1 ≤ *i* ≤ *k*, a lineage alive at 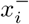 has no sampled extant descendant with probability 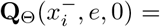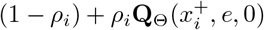 if *i* < *k* (since it is either not sampled at *x_i_* or sampled with no sampled extant descendant) and with probability 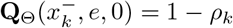 otherwise [32]. It follows that for all *t* ∈ [*x*_*i*−1_, *x_i_*], we have that

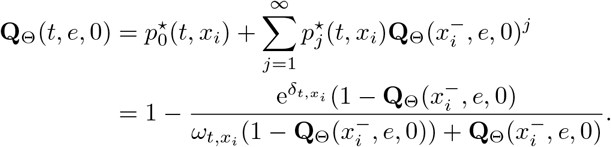

If *t* = *x*_*i*−1_, the formula above gives 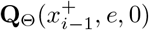, from which we get 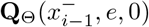 and iteratively any **Q**_Θ_(*t, e*, 0) from 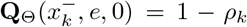. The probability **O**_Θ_(*t*) that a lineage alive at time *t* has at least a sampled extant descendant is basically 1 − **Q**_Θ_(*t, e*, 0) and can therefore be computed under the sampled-generalized-birth-death model Θ = (*s, e, λ, μ*, (*x_i_*, *ρ_i_*)_1≤*i*≤*k*_).

Let us compute **Q**_Θ_(*t, e*, 1), the probability that a lineage alive at *t* has exactly one sampled descendant at the end/present time *e*. For all 1 ≤ *i* ≤ *k* and all *t* ∈ [*x*_*i*−1_, *x_i_*], the probability that a lineage alive at *t* has a single lineage sampled at *e* is the sum over all *j* ≥ 1 of the probabilities that this lineage has *j* descendants at time 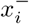 among which

- one is both sampled at *x_i_* and with a single descendant sampled at *e* and,
- *j* − 1 ones have no sampled descendants at *e*.

Namely, we have that

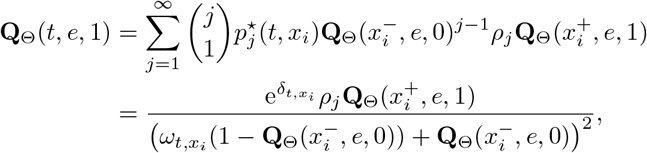

Since 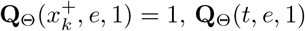 can be computed for all *t* ∈ [*s, e*] by iterating the formula above.

In order to show how to compute **Q**_Θ_(*s, e, N*), we shall first show that the probability density of observing the set of divergence times 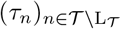 of any tree topology 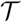 with *N* tips (i.e., with 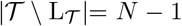 inner nodes and divergence times) under a sampled-generalized-birth-death model and disregarding the tree topology 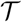 is

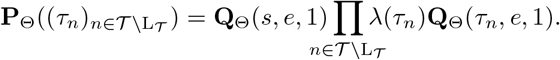

This result was actually already proved in [34, pp. 57-58]. But, since it was under the simple birth-death process and without including the first divergence time, let us sketch its proof.

The author of [34] first remarks that for all times *t* < *t′*, we have that

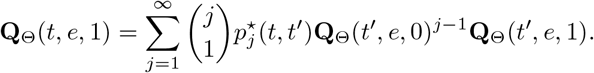

The equation above implies that the probability of observing no speciation event which gives rise to a lineage sampled at *e* between times *t* and *t′* on a lineage of the reconstructed tree is 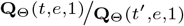.

For all internal nodes *n*, the branch ending by *n* ends at time *τ_n_* and starts at time 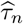 where 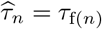 if *n* is not the root and f(*n*) is its direct ancestor, and where 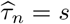 if *n* is the root. Considering the diversification process only on this branch, we observe a single lineage alive at 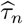 which goes to time *τ_n_* without (observable) speciation, which has probability 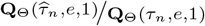. If *n* is a tip (i.e., 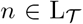) then we set *τ_n_* = *e* and we have **Q**_Θ_(*τ_n_, e*, 1) = 1. If *n* is an inner node (i.e., 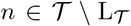) then we observe a speciation event at *τ_n_*, which occurs at rate *λ*(*τ_n_*). From the Markov property, all the branchs evolve independently and we get that the probability density of observing the divergence times 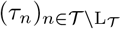 is

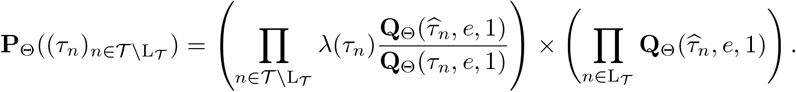

In the product above, each divergence time *τ_n_* occurs twice in numerators as 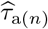 and 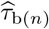, where a(*n*) and b(*n*) are the direct descendants of *n*, and once in denominators. The time origin occurs only once in the numerator associated to the root. By simplifying the product above, we eventually get that

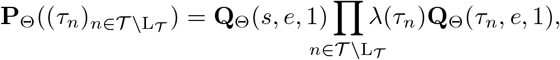

which does not depend on the tree topology 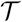 (except on its size) and which is the probability density of observing the divergence times 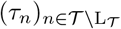 on any tree topology with *N* tips.

The probability of observing *N* lineages at *e* by starting with a single lineage at *s* (in any tree topology) is then obtained by integrating the probability density of *N* − 1 divergence times (*τ_j_*)_1≤*j*≤*N*−1_ between *s* and *e*:

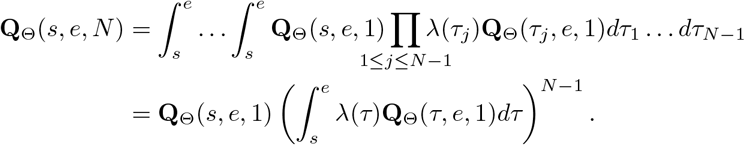

For all times *t* and *t′* with *s* ≤ *t* ≤ *t*′ ≤ *e*, the probability **Q**_Θ_(*t, t′, N*) can be computed in the same way by considering the restriction of the model Θ to the time interval [*t, t′*] (with full sampling at *t′* if *t′* ≠ *x_i_* for all 1 ≤ *i* ≤ *k*).

The probabilities of ending configurations of standard and special patterns under the sampled-generalized-birth-death model Θ = (*s, e, λ, μ*, (*x_i_*, *ρ_i_*)_1≤*i*≤*k*_) can then be computed from the probabilities **Q**_Θ_(*t, t′, N*) and **O**_Θ_(*t*) (Eq. 1 and 2). By putting Θ_[*t,t′*]_ for the model Θ restricted to the time interval [*t, t′*], we have that

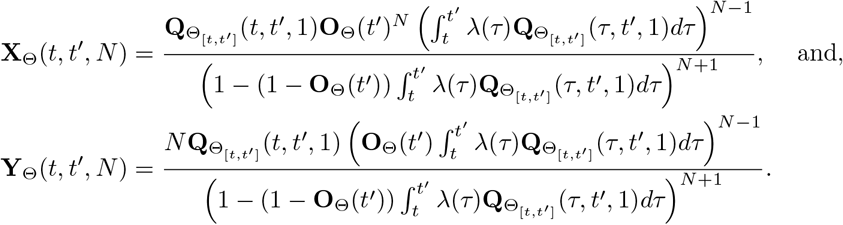

## C Proofs of Theorems

## C.1 Proof of Theorem 2

Let us start with the case where the oldest time is the end time of the diversification process, i.e., the case where *o* = *e*. By construction, we then have that 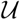 and 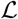 are both empty. It follows that 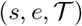 is a standard pattern of probability 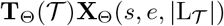 from Claim 1.

Let us now assume that *o* < *e*. Under the notations of the theorem and by assuming that the divergence times of 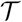 are consistent with the temporal constraints, let us define 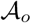 as the set of nodes of 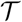 whose divergence times are anterior to *o* (i.e. 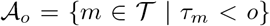). Since divergence times corresponding to ancestors of a given node are always posterior to its own divergence time, all sets 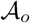 are start-sets. By construction, the set 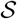 contains all the possible configurations of nodes of 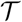 with divergence times anterior to *o* which are consistent with the temporal constraints 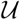 and 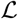. Since all these configurations are mutually exclusive, by putting 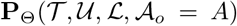 for the probability of observing the topology 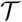 with 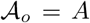 and the temporal constraints 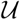 and 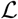, the law of total probabilities gives us that

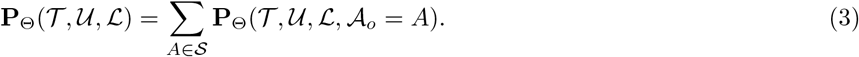

For instance, the entries of the second column of Figure 4 (just after the sum sign) represent all the start-sets of 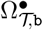.

In order to compute the probability 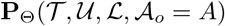 for a start-set 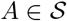, we remark that

- the part of the diversification process anterior to *o* is the standard pattern 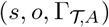 and that
- the part of the diversification process posterior to *o* consists of all the tree topologies 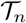 with temporal constraints 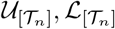 with 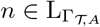 under the model Θ_[*o*]_ (i.e., the model Θ restricted to the interval of times [*o, e*]), which have probability 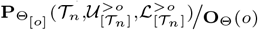 conditioned on the observability of their starting lineages.

Since the diversification model Θ is Markovian, evolution of all the tree topologies 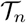 is independent of each other and with regard to the part of the process anterior to *o*, conditional upon starting with an observable lineage at time *o*.

From Claim 1, the probability of the standard pattern 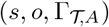 is 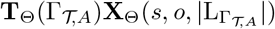 under the assumption that 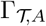 is labeled. This part is a little tricky since we don’t have a direct labeling of 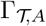 here (the tips of 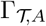 are identified through the labels of their tip descendants in 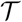, i.e., the tips of the subtrees pending from the tips of 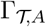). Since it assumes that 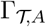 is (exactly) labeled, we have to multiply the probability obtained from Claim 1 with the number of ways of connecting the tips/labels of 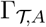 to the subtrees starting from *o*, which is 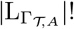, and with the probability of observing the groups of labels corresponding to the subtrees starting from *o*. Since all labelings of 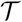 are equiprobable, the probability of the groups of labels corresponding to the subtrees starting from *o* is the inverse of the number of ways of choosing a subset of 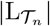 labels from 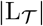 ones for all tips *n* of 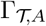 without replacement, i.e., the inverse of corresponding multinomial coefficient, which is

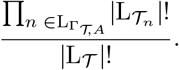

Putting all together, we eventually get that

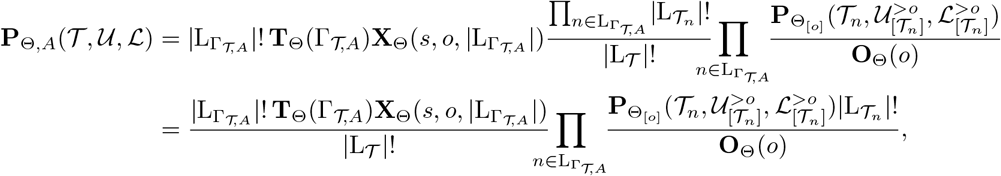

which, with Equation 3, ends the proof. The whole computation of a toy example is schematized in Figure 4.

## C.2 Proof of Theorem 3

Assuming that a diversification shift of the clade originating at *m* occurs at time *t* implies that the divergence times of 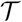 are such that both the direct ancestor of *m* has a divergence time strictly anterior to *t* and the divergence time of *m* is strictly posterior to *t*. Reciprocally, if the divergence times of *m* and of its direct ancestor are respectively posterior and anterior to *t*, then a diversification shift at time *t* may occur for the clade originating at *m*. The set of subsets of internal nodes with divergence time anterior to *t* consistent with the assumptions of the Theorem is thus exactly 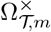.

We next follow the same outline as that of the proof of Theorem 2. For all subsets *A* of internal nodes of 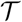, let us put 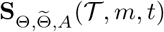 for the probability of observing the topology 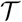 with a shift at time *t* for the clade originating at *m* and whose set of nodes with divergence time anterior to *t* is exactly *A*. We have that

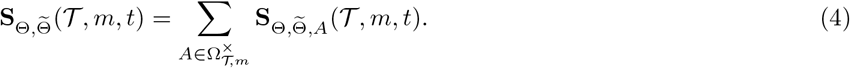

From the Markov property, we have that 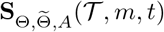 can be written as the product of the part of the diversification anterior to *t*, which is the special pattern 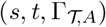 where the special lineage is the one on which the shift occurs, and the part of the diversification posterior to *t* which is a set of trees starting from time *t* and ending at time *e* by following model Θ_[*t*]_ except the special one which follows 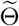. By construction, the non-special trees starting from *t* are conditioned on the observability of their starting lineage at *t*, thus have probability 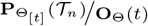 while the special one is not conditioned and has probability 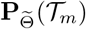.

From Claim 2, the probability of the special pattern 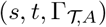 is 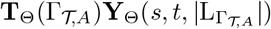 under the assumption that 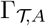 is labeled. The situation slightly differs from the case of a standard pattern treated in the proof of Theorem 2 since the special tip of the special pattern is well identified and so is the subtree pending from it. In order to taking into account the fact that 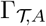 is not directly labeled, we have here to multiply the probability provided by Claim 2 with the number of ways of connecting the tips/labels of 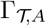 except *m*, the special one, to the subtrees starting from *t*, i.e., 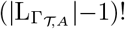, and with the probability of observing the groups of labels corresponding to the subtrees starting from *t*, which is

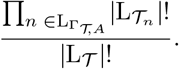

Eventually, we get that

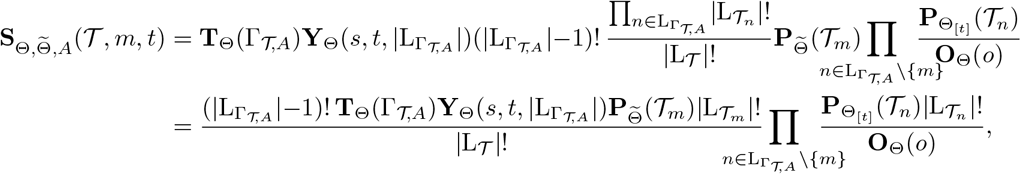

which with Equation 4 ends the proof.

## C.3 Proof of Theorem 4

If the model Θ is lineage homogeneous then, under the assumptions and notations of Theorem 2, we have that

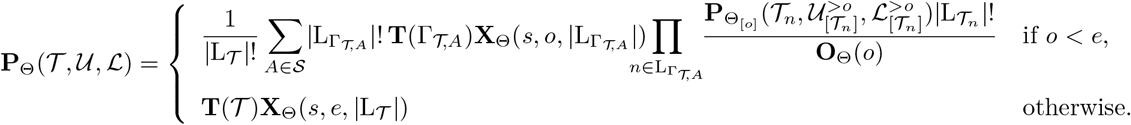

Since in the case where *o* = *e*, the computation of 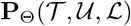 is performed in constant time under the assumptions of the theorem, we focus on the case where *o* < *e*. Let us first introduce an additional notation. For all sets 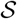 of start-sets of a tree topology 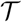 and all numbers *k* between 1 and the number of tips of 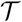, we put 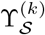 for the set of start-sets 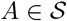 such that the corresponding start-tree 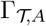 has exactly *k* tips. By construction, a start-tree of 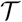 has at least one tip and at most 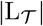 tips. We have:

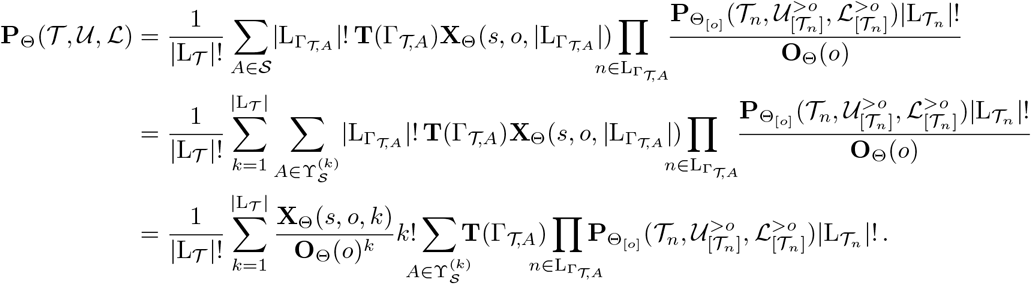

Let us set for all nodes *m* of 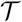,

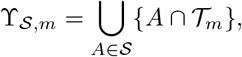

where 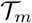 stands here for the set of nodes of the subtree topology rooted at *m*. In plain English, elements of 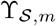 are elements of 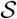 restricted to 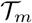. Since, by construction, the elements of 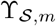 are start-sets of the tree topology 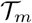, the start-tree 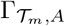 is well-defined for all 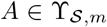. For all numbers 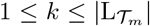, we put 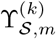 for the set of start-sets 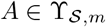 such that the corresponding start-tree 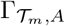 has exactly *k* tips.

Let us now define for all nodes *m* of 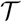 and all 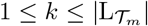, the quantity

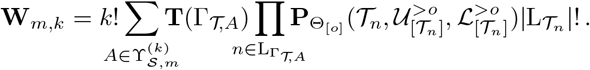

Basically, by putting *r* for the root of 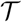, we have that

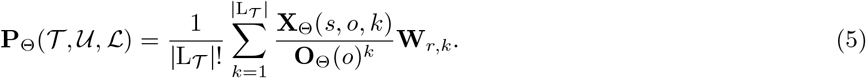

We shall see how to compute 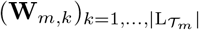 for all nodes *m* of 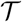.

Let us first consider the case where *k* = 1. We have that

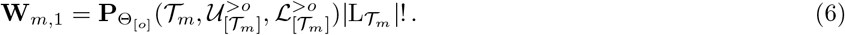

Let us now assume that *k* > 1 and let *a* and *b* be the two direct descendants of *m*. Since we assume *k* > 1, all start-sets of 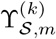 contain *m*. It follows that we have 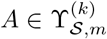 if and only if there exist two start-sets 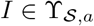 and 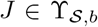 with {*m*} ∪ *I* ∪ *J* = *A*. The tree topology 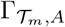 has root *m* with two child-subtrees 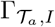 and 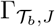. In particular, we have 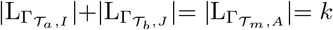.

From Theorem 1, we have that

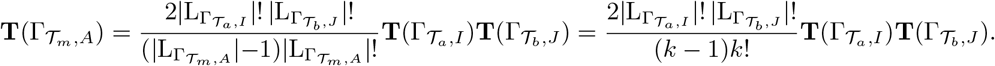

Moreover, since by construction 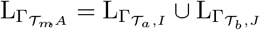, we get that

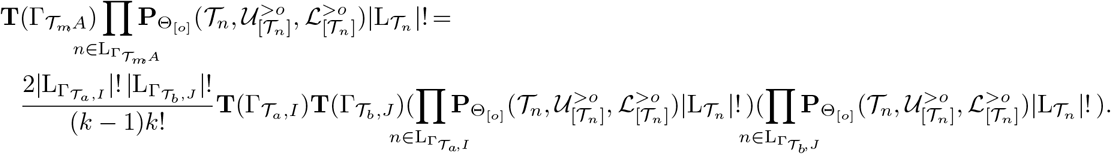

More generally, the start-sets of 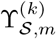 are in one-to-one correspondence with the set of pairs (*I, J*) of 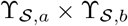 such that 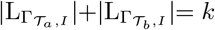. This set of pairs is exactly the union over all pairs of positive numbers (*i*, *j*) such that *i* + *j* = *k*, of the product sets of 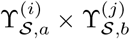. It follows that

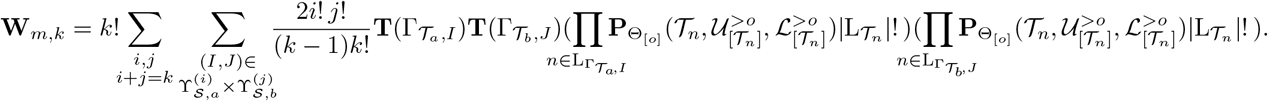

After factorizing the left hand side of the equation just above, we eventually get that for all *k* > 1,

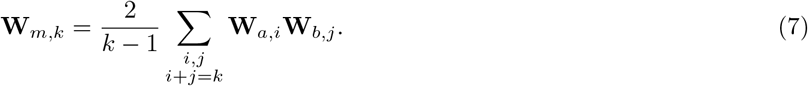

The following remark is straightforward to prove by induction.

### Remark 2.

*Let* 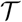 *be a binary tree topology and for all internal nodes n of* 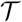, *let* a(*n*) *and* b(*n*) *denote the two direct descendants of n. We have that*

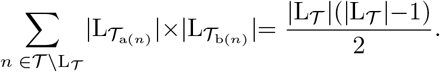

From Equation 7 and for all internal nodes *m* of 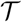 with children *a* and *b*, computing the quantities **W**_*m,k*_ for all 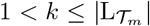 involves exactly 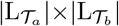 terms of the form **W**_*a,i*_**W**_*b,j*_. It follows that Remark 2 implies that if the quantities **W**_*m*,1_ are given for all nodes *m* of 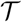, the quantities **W**_*m,k*_ for all 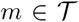 and all 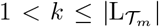 can be recursively computed in a time proportional to 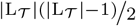, thus with time complexity 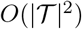.

In order to finish to prove Theorem 4, we remark that if 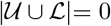 then 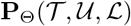 is the probability of the standard pattern 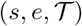 which can be computed with time and memory complexity *O*(1) from Claim 1.

In the case where 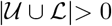, we shall proceed by induction on the total number temporal constraints by showing that the total time complexity required to compute the probabilities 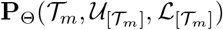 for all internal nodes *m* is 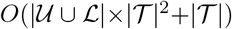.

This property is basically true in the base case where 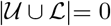, the probability 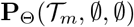 is the probability of a standard pattern which can be computed in constant time for all internal nodes *m* from Claim 1.

Let us assume that the induction assumption holds for all numbers of temporal constraints smaller than *ℓ* and let us consider two sets of temporal constraints 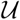 and 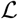 such that 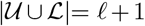. If *o* is the oldest time involved in 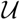 and 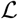, this implies that 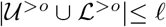. From the induction hypothesis, computing the probabilities 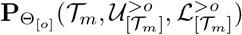 for all internal nodes *m* can be performed in 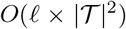. From Equation 6, the quantities **W**_*m*,1_ for all internal nodes *m* are calculated directly from the probabilities 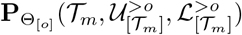, thus in 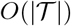. From Remark 2, all the quantities **W**_*m,k*_ for all internal nodes *m* of 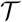 and all 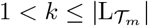 can be calculated with time complexity 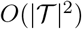. Equation 5 can then be applied to all subtrees of 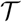 in order to compute the probabilities 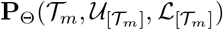 from the quantities **W**_*m,k*_ for all internal nodes *m* of 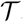. Since computing each 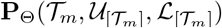 requires to sum 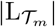 terms, computing all the 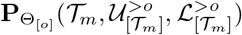 has total time complexity 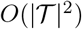.

To sum up, being given the probabilities 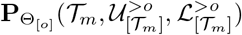, which can be computed with complexity 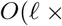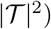 from the induction hypothesis, computing the probabilities 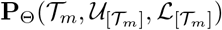 for all internal nodes *m* of 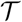 has time complexity 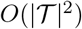. The total time complexity required to compute all the 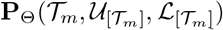 is well 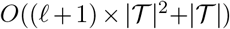. Since we assume that 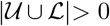, we have that 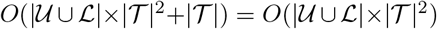, which ends to prove the statement of the theorem about the time complexity in this case.

Last, since at each stage of the induction, we have to store only the quantities 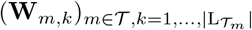 and the probabilities 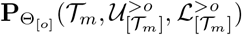 and 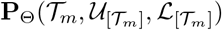 for all internal nodes *m* of 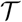, the total memory space complexity of the computation is 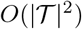.

## Notes

#### Summary of Updates

This revision contains minor changes and the presentation of an alternative method proposed by Amaury Lambert while reviewing the manuscript for PCIEvolBiol.

